# Asymmetric cell division safeguards memory CD8 T cell development

**DOI:** 10.1101/2022.11.08.515624

**Authors:** Fabienne Gräbnitz, Dominique Stark, Danielle Shlesinger, Anthony Petkidis, Mariana Borsa, Alexander Yermanos, Andreas Carr, Niculò Barandun, Arne Wehling, Miroslav Balaz, Timm Schroeder, Annette Oxenius

**Affiliations:** Institute of Microbiology, ETH Zurich, Vladimir-Prelog-Weg 4, 8093 Zurich, Switzerland.; Department of Biosystems Science and Engineering, ETH Zurich, Mattenstrasse 26, 4058 Basel, Switzerland.; Department of Molecular Life Sciences, University of Zurich, Winterthurerstrasse 190, 8057 Zurich, Switzerland.; The Kennedy Institute of Rheumatology, NDORMS, University of Oxford, Roosevelt Drive, Oxford OX3 7FY, UK; Center for Translational Immunology, University Medical Center Utrecht, Utrecht, Netherlands; Department of Metabolic Disease Research, Biomedical Research Center of the Slovak Academy of Sciences, Dubravska cesta 9, 845 05 Bratislava, Slovakia.; Department of Health Sciences and Technology, ETH Zurich, Schorenstrasse 16, 8603 Schwerzenbach, Switzerland.

## Abstract

The strength of T cell receptor (TCR) stimulation and asymmetric distribution of fate determinants are both implied to affect T cell differentiation. Here, we uncovered asymmetric cell division (ACD) as a safeguard mechanism for memory CD8 T cell generation specifically upon strong TCR stimulation. Using live imaging approaches, we found that strong TCR stimulation induced elevated ACD rates and subsequent single cell derived colonies comprised both effector and memory precursor cells. The abundance of memory precursor cells emerging from a single activated T cell positively correlated with first mitosis ACD. Accordingly, preventing ACD by inhibition of PKCζ during the first mitosis upon strong TCR stimulation markedly curtailed the formation of memory precursor cells. Conversely, no effect of ACD on fate commitment was observed upon weak TCR stimulation. Our data provide new mechanistic insights into the role of ACD for CD8 T cell fate regulation upon different activation conditions.

## Introduction

Robust and heterogeneous antigen-specific CD8 T cell responses are essential for effective host defense against infection. Upon initial acute viral infection, activated naïve CD8 T cells start to clonally expand and differentiate. The majority of generated CD8 T cells within this phase are effector cells (T_E_), responsible for rapid elimination of infected cells and viral clearance [1]. However, also CD8 memory precursor cells develop early during the clonal expansion phase at low frequencies [2]–[4]. While T_E_ are short-lived, leading to contraction of the overall response, memory CD8 T cells establish a long-lived pool composed of multiple distinct self-renewing subsets, which vary in their migratory and functional properties. Reactivation of these memory cells upon secondary encounter with the same pathogen leads to execution of immediate effector functions alongside secondary expansion, thereby providing protection [5]. Intriguingly, elegant studies have demonstrated that a single, naïve CD8 T cell can give rise to both effector and memory cells upon activation [6]–[9]. However, it remains unknown how divergence between effector and memory fates from a single naïve CD8 T cell is achieved on a mechanistic level. Different models propose various factors or mechanisms that contribute to the establishment of cellular heterogeneity, including the strength of initial T cell receptor (TCR) signaling and asymmetric cell division (ACD) [1], [10]. In both models, the immunological synapse (IS), formed between the antigen-presenting cell (APC) and the engaged T cell, orchestrates TCR activation and establishes a polarization axis serving as a prerequisite for ACD. TCR signal strength is impacted by TCR affinity towards its cognate antigen as well as antigen abundance and is suggested to modulate the proportional formation of effector and memory CD8 T cells, with strong TCR stimulation preferentially inducing short-lived effector cells (SLECs) and weaker stimulation favoring differentiation of memory precursor cells [11], [12]. In addition, it has been shown that strong TCR stimulation induces higher frequencies of cells undergoing ACD compared to weak TCR stimulation [11]. ACD is characterized by a polarized distribution of specific fate-determining transcription factors (e.g. Tbet and c-Myc), cell organelles (e.g. proteasomes), and surface receptors (e.g. CD8, TCR, CD25, IFNγR, Glut1). Further, it has been reported that the IS-proximal daughter cell - interacting with the APC - inherits molecules or organelles related to an effector fate, such as Tbet and CD25, whereas the IS-distal daughter cell inherits components promoting memory fate, such as the proteasome and PKCζ [10], [11], [13]–[21]. Previous studies have demonstrated a remarkable impact of differential expression of fate-determining markers on future fate of first daughter cells by using bulk sorting of cells either expressing the marker of interest at a high or low level after the first cell division upon activation, followed by downstream analyses investigating their future fate [20]–[23]. Furthermore, the role of ACD in fate diversification is supported by the reported transcriptional heterogeneity within CD8 T cells that have undergone one cell division after activation [22], [24]. However, a direct link of ACD to subsequent asymmetric fate on a single cell level is missing. Therefore, tracing of individual daughter cells that emerge from an ACD and analysis of their progeny with respect to their differentiation is necessary [25]. As various studies provide evidence for a significant impact on fate diversification for both models - ACD and TCR signal strength - we aimed to elucidate their interplay. To this end, we established experimental systems allowing to follow the fate of single daughter cell progenies derived from an ACD either induced by weak or strong TCR stimulation using live imaging. We validated and used the combinatorial expression of T cell factor 1 (TCF1) and L-selectin (CD62L) as markers indicating effector and memory fate specification a few days after activation. Using these two markers as predictors of future fate in combination with long-term live imaging approaches, we found that strong TCR stimulation led to elevated ACD rates and single cells undergoing ACD established mixed-fate colonies comprising both effector and memory precursor cells. Strikingly, experimental impairment of ACD during the first cell division after activation upon strong TCR stimulation markedly curtailed the development of memory precursor cells, which resulted in limited memory formation *in vitro* and *in vivo*. In contrast, upon weak TCR stimulation, ACD was not associated with different cell fates and single activated cells formed exclusively single-fate colonies, either comprising memory or effector precursor cells, irrespectively of the asymmetry of their first cell division. In addition, no effect of ACD inhibition was observed on the formation of memory precursor cells. Together, our results indicate that ACD during the first mitosis after activation functions as a safeguard mechanism for CD8 T cell memory formation following strong TCR stimulation.

## Results

### Expression of TCF1 and CD62L identifies early memory and effector precursor CD8 T cells

To relate divergent cell fates of activated CD8 T cells to ACD, we established an *in vitro* CD8 T cell activation protocol that allows tracking of individual cells and their progeny over several generations by live microscopy. Monitoring of early differentiation states indicative of memory or effector cell differentiation requires the identification of early fate determination markers. Previous studies have shown that CD62L and TCF1 are expressed in naïve and memory CD8 T cells and are both downregulated in effector cells [26], [27]. Furthermore, CD62L^+^TCF1^+^ cells generated *in vivo* during the early acute effector phase were shown to give rise to central memory T cells (T_CM_) [2], [3]. We therefore investigated whether *in vitro* stimulation of CD8 T cells also gives rise to an early establishment of CD62L^+^TCF1^+^ cells alongside CD62L^-^TCF1^-^ cells, and whether expression of TCF1 and CD62L at these early stages of differentiation serve as reliable markers indicating future fate. To this end, we used TCR transgenic P14 CD8 TCF1-GFP cells, which specifically recognize the gp_33-41_ peptide from the Lymphocytic Choriomeningitis Virus (LCMV) glycoprotein and additionally express a *Tcf7^GFP^* reporter [28]. P14 CD8 TCF1-GFP cells were activated by plate bound α-CD3 and α-CD28 antibodies in addition to Fc-ICAM-1 and IL-2 for 36-40 hours (h), then removed from the activation stimuli and further cultured in the presence of IL-2, IL-7 and IL-15 (Fig. 1A). 4 days later, we observed marked proliferation and partial downregulation of TCF1 and CD62L, resulting in the development of three populations identified by divergent expression of TCF1 and CD62L (CD62L^+^TCF1^+^, CD62L^+^TCF1^-^, CD62L^-^TCF1^-^, Fig. 1A and 1B). CD62L required at least 4 cell divisions before downregulation was initiated, while TCF1 downregulation started after 2 - 3 cell divisions (Fig. 1C). Next, we dissected the *in vitro* behavior of these three populations. In line with the finding that both markers required several rounds of cell division to initiate downregulation, we found that CD62L^-^TCF1^-^ cells proliferated the most, whereas CD62L^+^TCF1^-^ cells were slower and CD62L^+^TCF1^+^ cells underwent the least rounds of cell division on day 4 and day 7 after activation (Fig. 1D). Furthermore, sorting and subsequent individual culture of the three subsets revealed that the CD62L^+^TCF1^+^ subset partially fed into the two other subsets, while CD62L^+^TCF1^-^ cells only fed into the CD62L^-^TCF1^-^ subset and the CD62L^-^TCF1^-^ subset mainly preserved its phenotype 2 days after additional culture, indicating a clear direction of differentiation (Fig. 1E). In further analyses, we focused on the most distinct CD62L^+^TCF1^+^ and CD62L^-^TCF1^-^ cell subsets and wondered whether the observed differences in proliferation and resulting expression of CD62L and TCF1 might be associated with distinct expression profiles of costimulatory and coinhibitory receptors. In line with previous data, showing that early memory precursor cells are maintained by inhibitory signaling [2], CD62L^+^TCF1^+^ cells expressed elevated levels of PD1. Instead, CD62L^-^TCF1^-^ cells showed higher expression of CD25, providing enhanced sensitivity to proliferation inducing IL-2 (Fig. 1F). Additionally, TIM3 and ICOS were found to be higher expressed in CD62L^-^TCF1^-^ cells compared to CD62L^+^TCF1^+^ cells (Fig. S1A and B), whereas for TIGIT, TOX, CTLA-4 and Lag3 no differences were observed (data not shown). The transcription factor FOXO1 was previously described to induce expression of CD62L and TCF1 in addition to other memory specific markers [29]–[31]. Using confocal microscopy, we investigated the cellular localization of FOXO1 and found that in naïve CD8 T cells and in early establishing CD62L^+^TCF1^+^ CD8 T cells upon activation, FOXO1 was mainly localized in the nucleus, enabling transcription of CD62L and TCF1, while in CD62L^-^TCF1^-^ cells, FOXO1 was mainly found in the cytoplasm (Fig. 1G). We next characterized the functional profiles of CD62L^+^TCF1^+^ and CD62L^-^TCF1^-^ cells established early after activation. Memory and effector CD8 T cells differ substantially in their metabolic profiles. While naïve and memory CD8 T cells primarily rely on oxidative phosphorylation, effector CD8 T cells rely mainly on glycolysis [31]–[33]. To investigate the metabolic profiles of the two subsets, we performed extracellular flux analysis. Interestingly, CD62L^+^TCF1^+^ cells demonstrated higher basal mitochondrial respiration and ATP production compared to CD62L^-^TCF1^-^ cells, while CD62L^-^TCF1^-^ cells had higher extracellular acidification rates (ECAR) over time and significantly higher glycolysis and glycolytic capacity (Fig. 1H). Re-stimulation with gp_33-41_ of CD62L^+^TCF1^+^ cells resulted in pronounced IL-2 production, whereas CD62L^-^TCF1^-^ cells produced more IFNγ and TNFα compared to CD62L^+^TCF1^+^ cells (Fig. 1I). These results demonstrate that *in vitro* generated CD62L^+^TCF1^+^ cells possess characteristics of memory cells, whereas CD62L^-^TCF1^-^ cells are endowed with effector cell features.

**Figure 1:**
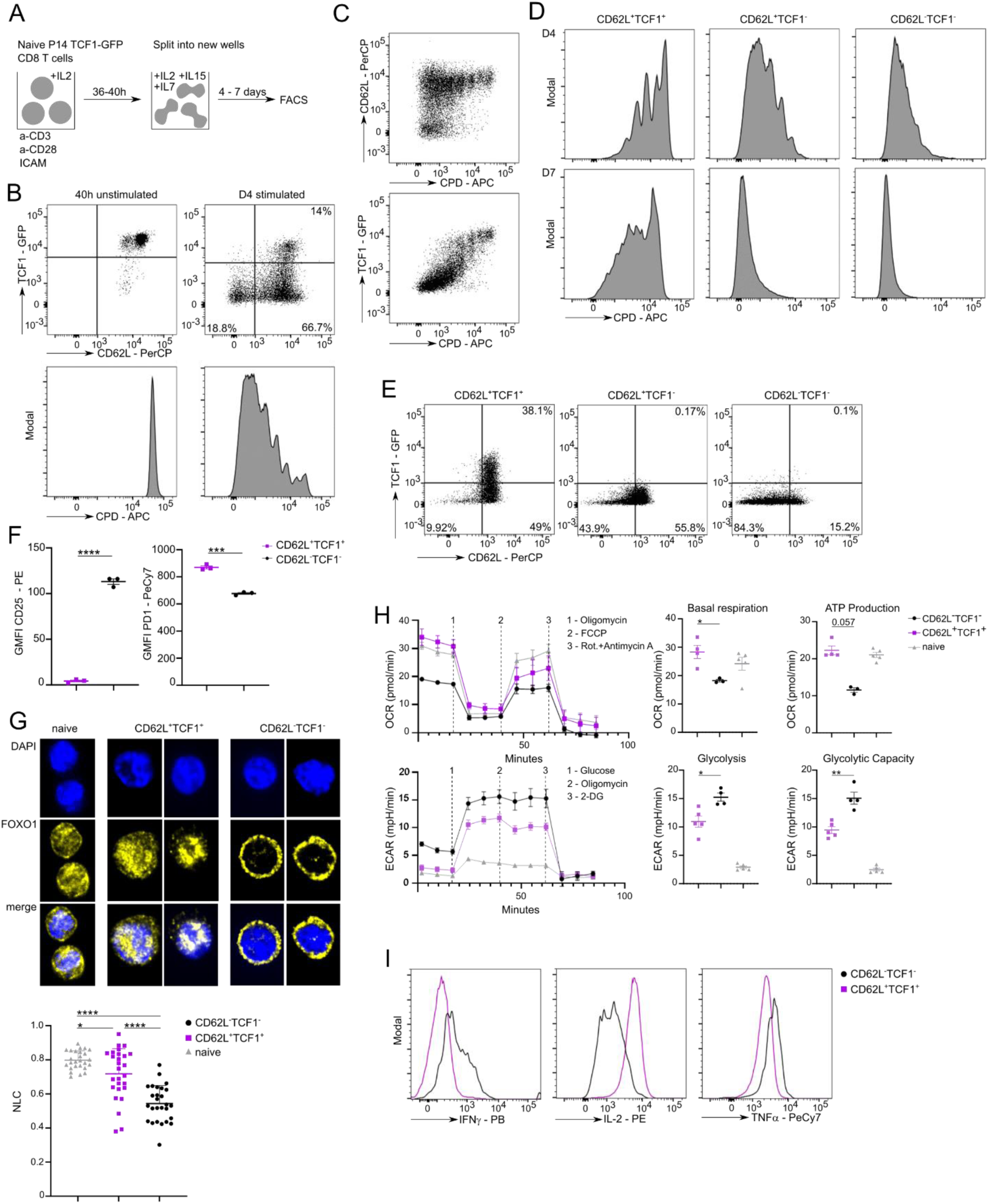
*In vitro* differentiation of P14 TCF1-GFP cells and characterization of CD62L^+^TCF1^+^ and CD62L^-^TCF1^-^ cells. **A** Experimental setup. Naïve P14 TCF1-GFP cells were stimulated with plate-bound α-CD3 and α-CD28 antibodies and Fc-ICAM-1 in the presence of IL-2 for 36 – 40 h before cells were transferred to new wells in medium containing IL-2, IL-7 and IL-15. Cells were cultured for 4 – 7 days until FACS analysis. **B** Representative FACS plots of TCF1 and CD62L expression in P14 TCF1-GFP cells and respective cell proliferation dye (CPD) dilution with or without stimulation after 4 days. **C** Representative FACS plots of P14 TCF1-GFP cells expressing CD62L and TCF1 against CPD dilution on day 4 post stimulation. **D** Representative CPD dilution of CD62L^+^TCF1^+^, CD62L^+^TCF1^-^ and CD62L^-^TCF1^-^ cells on day 4 and 7 post stimulation. **E** Representative FACS plots of TCF1 and CD62L expression. P14 TCF1-GFP cells were stimulated for 4 days and sorted into CD62L^+^TCF1^+^, CD62L^+^TCF1^-^ and CD62L^-^TCF1^-^ cells. Subsets were individually re-cultured for 2 days before CD62L and TCF1 expression was re-assessed. **F** Geometric Mean of Fluorescence Intensity (GMFI) of CD62L^+^TCF1^+^ and CD62L^-^TCF1^-^ P14 TCF1-GFP cells expressing CD25 and PD1 on day 4 post stimulation. **G** Cellular localization of FOXO1. Representative confocal microscope images of fixed naïve (n=26), CD62L^+^TCF1^+^ (n=27) and CD62L^-^TCF1^-^ (n=27) cells. Distribution of nuclear localization coefficient (NLC) in all three cell subsets. **H** Oxygen consumption rate (OCR) (upper panel) and extracellular acidification rate (ECAR) (lower panel) of naïve, d5 sorted CD62L^+^TCF1^+^ and CD62L^-^TCF1^-^ cells was measured under basal conditions and in response to indicated drugs. **I** Representative histograms depicting production of IFNγ, IL-2 and TNFα by CD62L^+^TCF1^+^ and CD62L^-^TCF1^-^ cells. P14 cells were stimulated and sorted on day 4 into the two subsets followed by re-stimulation with gp33-41 for 6 h at 37°C. (B to D) Representative data from one of four experiments. (H) Representative data from one of two experiments. (E) Representative data from one of two experiments. (F) Representative data from one of two experiments. (I) Representative data from one of three experiments. Statistical analysis was performed using the unpaired two-tailed Student’s *t* test or, when data did not pass the Shapiro-Wilk normality test, the unpaired two-tailed Mann-Whitney test. \**P* < 0.05; \*\**P* < 0.01; \*\*\**P* < 0.001; \*\*\*\**P* < 0.0001.

### Adoptively transferred CD62L^+^TCF1^+^ cells home better to lymphoid organs and give rise to memory cells upon LCMV challenge

As *in vitro* generated CD62L^+^TCF1^+^ cells were characterized by memory features and CD62L^-^TCF1^-^ cells by effector hallmarks, we next addressed the *in vivo* behavior of the two subsets upon adoptive transfer. To this end, P14 TCF1-GFP cells were activated as described in Figure 1A, sorted on day 4 post activation into CD62L^+^TCF1^+^ and CD62L^-^TCF1^-^ cells and adoptively transferred into naïve recipient B6 mice at equal numbers. First, we assessed homing to lymphoid and peripheral organs 8h post cell transfer (Fig. 2A). While significantly more CD62L^+^TCF1^+^ cells localized to the lymph nodes compared to CD62L^-^TCF1^-^ cells, no significant differences were observed in the abundance of the two subsets in the spleen and lung (Fig. 2B). Furthermore, we prepared fixed spleen slices and analyzed *in situ* localization of transferred P14 cell subsets by confocal microscopy. CD62L^+^TCF1^+^ cells entered the T cell zone at significantly higher numbers compared to CD62L^-^TCF1^-^ cells, which were positioned mostly outside of the T cell zones (Fig. 2C). These data indicate that CD62L^+^TCF1^+^ cells have an advantage in homing to the lymph nodes, specifically to the T cell zones, early after adoptive transfer. We next determined the re-expansion potential and differentiation into secondary effector and memory cells of the two subsets after adoptive transfer followed by acute LCMV WE challenge one day later (Fig. 2D). 31 days post infection, the offspring of CD62L^+^TCF1^+^ cells were found at significantly higher numbers in lymph nodes and spleens of recipient mice. Furthermore, CD62L^+^TCF1^+^ cells gave rise to significantly higher numbers of memory cells (IL7R^+^KLRG1^-^) in lymph nodes and spleens (Fig. 2E). This trend was also observed in the lungs. We observed altered expression of CD62L and TCF1 31 days post infection when compared to the phenotype on the day of initial transfer (Fig. 2F). While the predominant phenotype of progeny cells coming from both subsets in the lung and spleen of recipient mice was CD62L^-^TCF1^-^, the phenotype of cells in the lymph nodes was more heterogeneous with mainly CD62L^+^TCF1^+^ and CD62L^-^TCF1^+^ cells coming from the CD62L^+^TCF1^+^ group, while the less abundant progeny of the CD62L^-^TCF1^-^ group mainly comprised cells with a CD62L^-^TCF1^+^ and CD62L^-^TCF1^-^ phenotype (Fig. 2F). Altogether, these findings indicate that *in vitro* generated CD62L^+^TCF1^+^ cells, as early as on day 4 post activation, can be characterized as memory precursor cells, whereas CD62L^-^TCF1^-^ cells at the same time can be classified as effector precursor cells.

**Figure 2:**
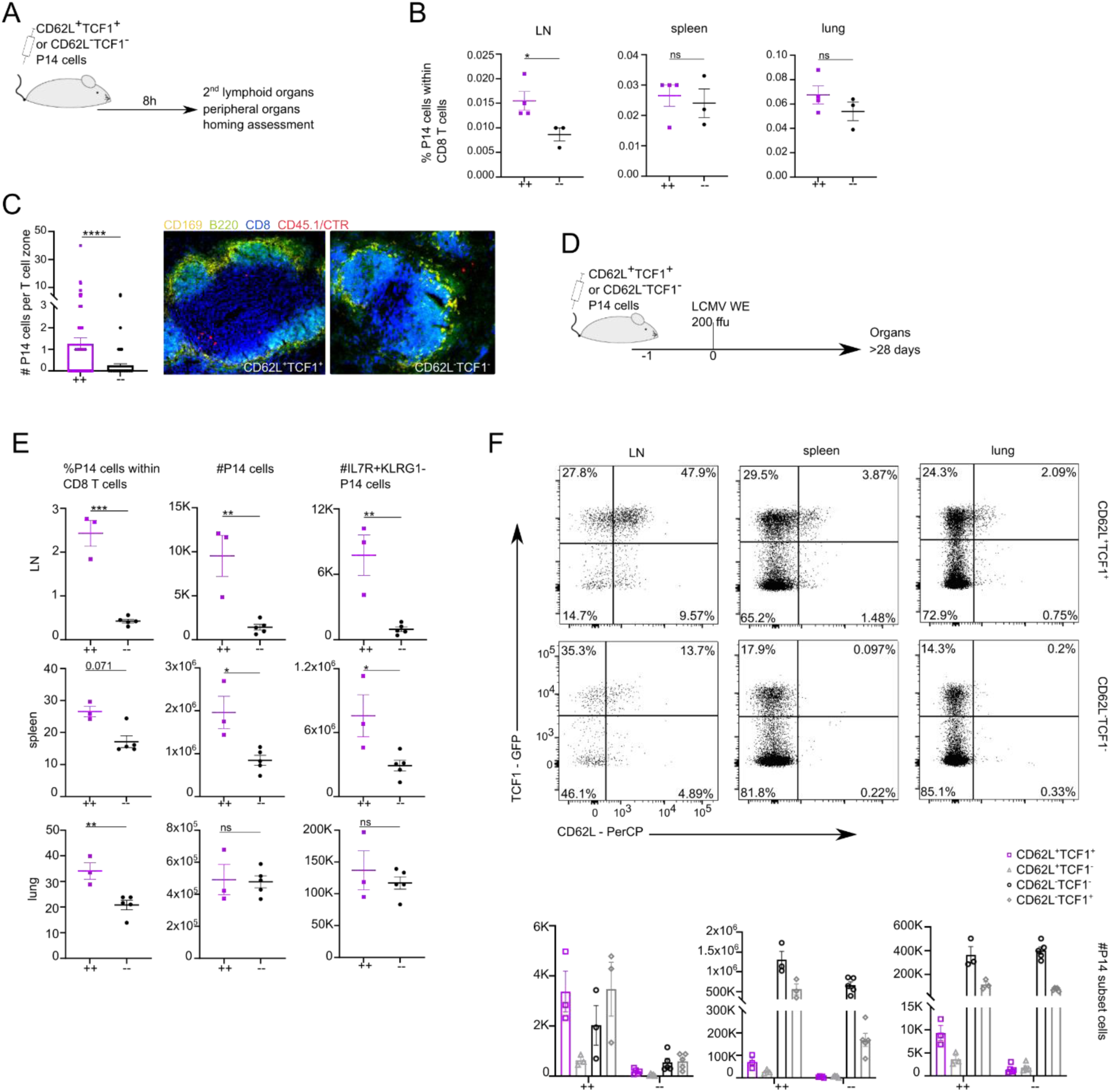
Adoptively transferred CD62L^+^TCF1^+^ cells home better to lymphoid organs and give rise to memory cells upon LCMV challenge. P14 TCF1-GFP cells were activated as described in Figure 1A and sorted on day 4 post activation into CD62L^+^TCF1^+^ and CD62L^-^TCF1^-^ cells. Sorted subsets were individually transferred at equal numbers into B6 recipient mice. **A** Experimental setup. Spleens, lymph nodes and lungs were harvested 8h post transfer and homing of transferred cells was investigated. **B** Frequencies and absolute numbers of P14 subset cells within spleen, lymph nodes and lung of recipient mice. **C** Left: Absolute numbers of splenic P14 cells localized in T cell zones. Right: Representative confocal microscopy images of 10-µm fixed splenic sections. Tissues were stained for the localization of metallophilic macrophages (CD169), B cells (B220), and CD8 T cells (CD8). P14 cells could be identified by CD45.1 and preserved CTR staining. **D** Experimental setup. Mice were infected with acute LCMV WE (200 ffu/mouse intravenously) one day after adoptive transfer. Spleens, lymph nodes and lungs were harvested 31 days post infection. **E** Frequencies and absolute numbers of P14 cells within spleens, lymph nodes and lungs of recipient mice. Absolute numbers of IL7R^+^KLRG1^-^ cells. **F** Representative FACS plots of TCF1-GFP and CD62L expressing P14 cells in lymph nodes, spleens and lungs of recipient mice. Absolute numbers of CD62L^+^TCF1^+^, CD62L^+^TCF1^-^, CD62L^-^TCF1^-^ and CD62L^-^TCF1^+^ cells. (A - C) Representative data from one of three experiments. (D - F) Representative data from one of two experiments. Statistical analysis was performed using the unpaired two-tailed Student’s *t* test or, when data did not pass the Shapiro-Wilk normality test, the unpaired two-tailed Mann-Whitney test. \**P* < 0.05; \*\**P* < 0.01; \*\*\**P* < 0.001; \*\*\*\**P* < 0.0001.

### ACD does not promote diversity upon stimulation with TCR agonistic antibodies

The identification of the two early fate indicating CD62L^+^TCF1^+^ and CD62L^-^TCF1^-^ cell subsets allowed us to investigate whether ACD might be a key determinant in effector and memory precursor cell generation originating from one naïve mother cell. To this end, we activated naïve P14 TCF1-GFP cells with plate-bound α-CD3, α-CD28, Fc-ICAM-1 and IL-2 for 32 – 34 h before the first cell division occurred (Fig. 3A and B). Such activation was sufficient to induce full activation of the cells, indicated by high CD44 expression (Fig. 3B). The subset of unstimulated CD44^-^ P14 cells (∼30%) present in the activation condition was neglected and excluded from downstream analyses, as CD44^-^ cells did not proliferate until day 4 post activation (Fig. S2A). After 32 – 34 h, we harvested the P14 cells and transferred them onto imaging slides in medium containing IL-2, IL-15 and IL-7, and started time-lapse imaging for 3 days (Fig. 3A) [34], [35]. To facilitate precise tracking [36], we established a coating with α-CD44 and α-CD43 antibodies, which was applied to the imaging slides prior to adding the cells. This coating led to adherence of the cells [37] and therefore allowed precise tracing of individual single cells without inducing further activation signals assessed by unaltered phospho-Akt (pAkt) staining, or inducing changes of the proliferation profile or alterations of expression of TCF1 and CD62L (Fig. 3C). We analyzed asymmetry between the daughter cells of first mitoses by differential expression of the surface marker CD8, which has been described as a reliable readout for asymmetric CD8 T cell division in previous studies [15], [22], [38]. Cell divisions were defined as asymmetric when the CD8 signal was 1.5-fold greater in one daughter cell compared to the other, corresponding to an asymmetry rate of 0.2. P14 cells divided asymmetrically as well as symmetrically in their first cell division (Fig. 3D). The frequencies of asymmetrically dividing cells were similar to those observed when ACD was measured in mitotic cells from fixed samples by confocal microscopy [22]. Throughout the time-lapse movie, we observed the occurrence of “big” (> 6 cell divisions) and “small” (2 - 5 cell divisions) colonies (Movie 1 and 2). Small colonies contained cells of comparably small individual cell size, while the cells within big colonies showed a larger cellular size (Fig. S2C). Furthermore, while cells giving rise to big colonies divided throughout the entire imaging period every 6 – 8 h, cells within small colonies stopped dividing after 3 - 4 cell divisions and entered a quiescent state (Fig. S2D). Continuous cell division of cells within big colonies prevented precise tracking after the 4^th^ - 5^th^ cell division as the cells were positioned around and on top of each other. We then investigated the phenotype of the evolving colonies and found that in line with the described proliferation profile (Fig. 1D), small colonies exclusively contained CD62L^+^TCF1^+^ cells, whereas big colonies were strictly composed of CD62L^-^TCF1^-^ cells (Fig. 3E). In accordance with the described “intermediate” population (Fig. 1D and E), we also observed colonies consisting of CD62L^+^TCF1^-^ cells. Interestingly, we did not observe any mixed-fate colonies, i.e. family trees that would comprise both cells with a CD62L^+^TCF1^+^ and CD62L^-^TCF1^-^ phenotype, which would be compatible with an ACD-induced bifurcation of fates. Instead, the evolving colonies stemming from one mother cell were strictly homogeneous in expression of CD62L and TCF1. We then correlated the CD8 (a)symmetry of the first cell division of each individual cell with fate outcome of their emerging progeny and found that differentiation into either an effector precursor or memory precursor population was independent of the degree of asymmetry of the first cell division (Fig. 3F). The frequency of first symmetric and asymmetric cell divisions was similar between the ensuing CD62L^-^TCF1^-^, CD62L^+^TCF1^-^ and CD62L^+^TCF1^+^ colonies, indicating no direct impact of ACD on fate determination under the applied stimulation conditions.

**Figure 3:**
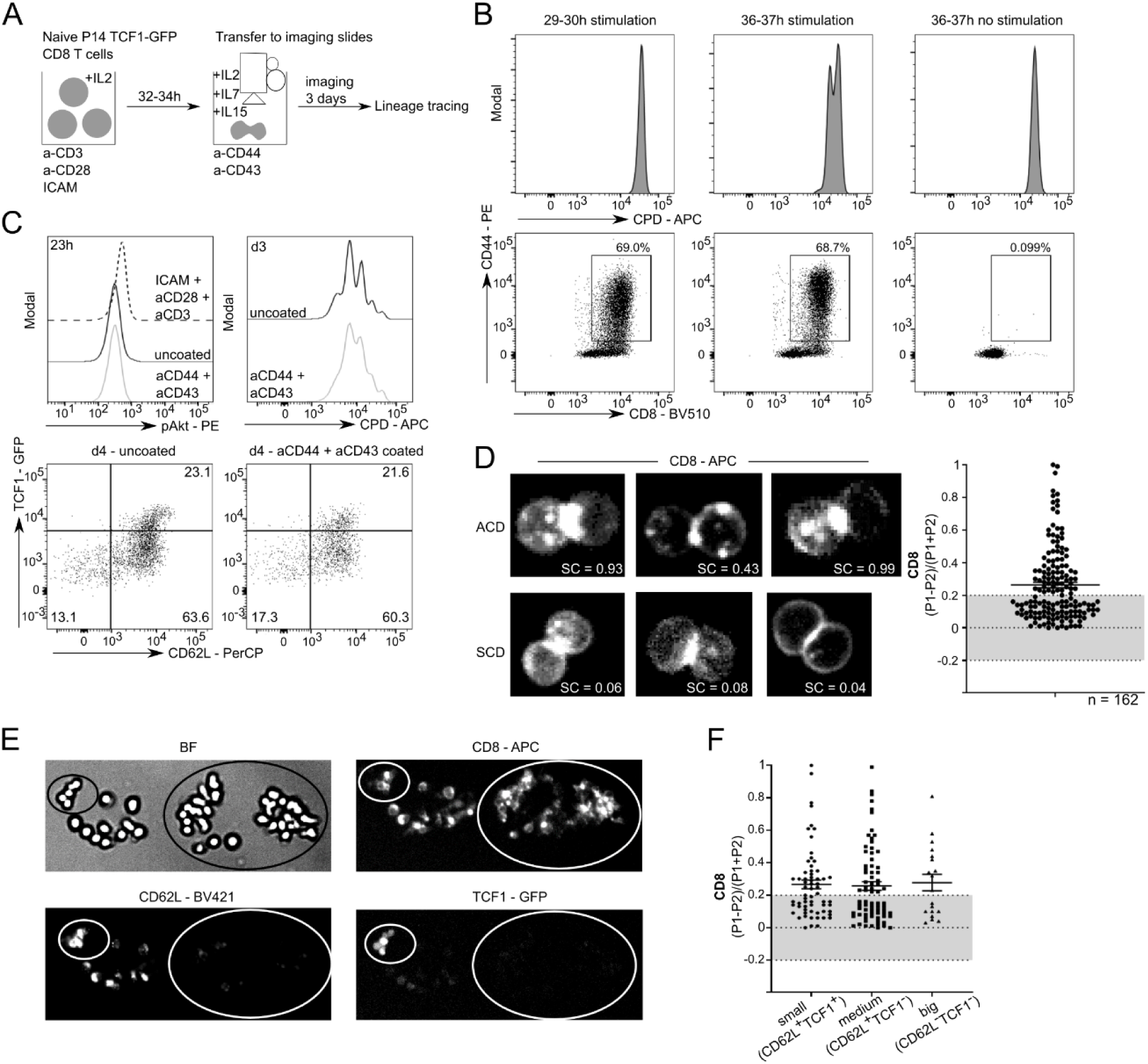
ACD does not promote diversity upon stimulation with TCR agonistic antibodies. **A** Experimental setup. P14 TCF1-GFP cells were activated as described in Figure 1A for 32 – 34 h. Cells were harvested and transferred onto imaging slides pre-coated with α-CD44 and α-CD43 antibodies in imaging medium containing IL-2, IL-7 and IL-15. Time-lapse imaging was performed for 3 days with 10× magnification. BF and far red channels were acquired every 40 min, blue and green channels were acquired every 90 min. **B** Representative plots of CPD dilution and CD44 expression of unstimulated and activated P14 TCF1-GFP cells after 29 – 30 h and 36 – 37 h. **C** P14 TCF1-GFP cells were activated as described in Figure 1A for 34 h and further cultured on uncoated or α-CD44 and α-CD43 antibody coated surfaces. Representative plots of pAkt (23 h post seeding), CPD dilution (day 3 post seeding) and TCF1 and CD62L expression (day 4 post seeding). As a positive control for pAkt staining, naïve P14 TCF1-GFP cells were stimulated on Fc-ICAM-1, α-CD3 and α-CD28 in the presence of IL-2. **D** Representative time-lapse images of asymmetric (ACD) and symmetric cell divisions (SCD) and quantified ACD rates based on CD8-APC surface expression. CD8 staining was quantified in both daughter cells and P1 was arbitrarily defined as the pole with higher amounts of CD8. Cell division was classified as asymmetric when the amount of CD8 was 50% higher in one daughter cell compared to the other one, defining the threshold of 0.2 (dashed line). Data are represented as mean ± SEM. E Representative time-lapse images of colonies on day 3 depicted in BF, CD8-APC (far red channel), CD62L-BV421 (blue channel) and TCF1-GFP (green channel). Circles are drawn around colonies stemming from one cell. F ACD rates from P14 TCF1-GFP cells that later formed small-sized (CD62L^+^TCF1^+^), medium-sized (CD62L^+^TCF1^-^) and big-sized (CD62L^-^TCF1^-^) colonies. Data are represented as mean ± SEM. (D to G) Data pooled from two independent experiments.

### The strength of TCR stimulation impacts ACD and fate

As ACD did not lead to different fates within the emerging progeny of single cells under the above applied stimulation conditions (i.e. α-CD3, α-CD28 and Fc-ICAM-1), we wondered whether varying the strength of initial TCR stimulation might have an impact on ACD promoting bifurcation of fates. TCR stimulation strength can be modulated by both TCR-pMHC affinity and pMHC density on antigen-presenting cells, leading to differences in the kinetics and magnitude of the CD8 T cell response [11], [39]–[44]. We decided to use two distinct peptide modalities of the LCMV glycoprotein, the variant C6 (KAVYNCATC), providing low peptide affinity for the P14 TCR, and the wild-type high affinity variant gp33 (KAVYNFATC) [45]. Moreover, we used the high affinity peptide variant gp33 at two concentrations (10^-6^ M and 10^-11^ M) to additionally probe for an effect of antigen abundance. The C6 variant was used at 10^-6^ M. To provide optimal CD8 T cell activation, we stimulated adherent dendritic cells (DCs) from the MutuDC1940 cell line [46] with CpG to induce expression of costimulatory molecules CD80, CD86 and CD40 (Fig. S3A), thus facilitating the establishment of an IS, and loaded the peptides onto their MHC molecules. Then, we added naïve P14 TCF1-GFP T cells for 28 – 30 h before we transferred them into new wells containing medium with IL-2, IL-7 and IL-15 for further culture (Fig. 4A). Compared to P14 CD8 T cells activated by plate-bound antibodies, pMHC/DC-activated P14 cells already performed their first cell division 28 – 30 h post activation (Fig. 4B). This was true for both peptide variants. We then investigated proliferation and phenotype of the P14 cells on day 4 or 6 post stimulation. Interestingly, while P14 cells activated by gp33 at 10^-6^ M proliferated the most, gp33 at 10^-11^ M and C6 stimulated P14 cells proliferated slower but in a comparable manner (Fig. 4C). Furthermore, gp33 at 10^-11^ M and C6 activation induced comparable frequencies of CD62L^+^TCF1^+^ and CD62L^-^TCF1^-^ cells whereas gp33 at 10^-6^ M induced significantly fewer CD62L^+^TCF1^+^ cells but more CD62L^-^TCF1^-^ cells (Fig. 4D and E). Overall, expression levels of activation markers such as CD25, PD-1 and ICOS were higher after gp33 10^-6^ M stimulation compared to gp33 10^-11^ M and C6 stimulation (Fig. S3B) and CD62L^-^TCF1^-^ cells expressed higher levels compared to CD62L^+^TCF1^+^ cells. Only after weak TCR stimulation, PD1 expression was found to be slightly higher on CD62L^+^TCF1^+^ cells compared to CD62L^-^TCF1^-^ cells. Comparing the phenotype and proliferation data of P14 cells between plate-bound antibody stimulation (Fig. 1) and pMHC/DC-activation, indicates that the outcome of plate-bound antibody stimulation resembles the two low TCR stimulation strength conditions, i.e., gp33 at 10^-11^ M and the C6 variant. We then assessed the emerging subpopulations regarding TCF1 and CD62L expression in more detail. Proliferation analyses indicated that CD62L^-^TCF1^-^ cells proliferated the most, whereas CD62L^+^TCF1^-^ cells cycled slightly slower and CD62L^+^TCF1^+^ cells showed the lowest number of cell divisions (Fig. 4F). To determine the behavior of CD62L^+^TCF1^+^ and CD62L^-^TCF1^-^ cells after *in vivo* stimulation, we sorted P14 TCF1-GFP cells 6 days after initial activation, provided either by the high or the low affinity peptide, into the two subsets, and adoptively transferred them into recipient mice followed by acute LCMV WE infection. Consistent with our previous findings, the frequencies and absolute numbers of P14 cells deriving from CD62L^+^TCF1^+^ cells were significantly higher in spleens and lymph nodes of recipient mice on day 35 post infection compared to CD62L^-^ TCF1^-^ derived P14 cells (Fig. S3C). The absolute number of P14 IL7R^+^KLRG1^-^ memory cells deriving from CD62L^+^TCF1^+^ cells was also significantly higher compared to CD62L^-^TCF1^-^ derived P14 cells in spleens and lymph nodes (Fig. S3C). These findings were observed for both peptide stimulations. TCF1 and CD62L expression of P14 cells in spleens and lymph nodes was comparable to P14 cells stimulated with TCR agonistic antibodies (Fig. S3C and Fig. 2F). To compare memory potential upon strong versus weak TCR stimulation on a bulk population level, we adoptively transferred P14 TCF1-GFP cells into recipient mice on day 6 after activation by pMHC/DC. Mice were challenged immediately with LCMV WE one day later. At day 35 post infection, frequencies and numbers of P14 cells derived from C6 and gp33 10^-11^ M were significantly higher in lymph nodes of recipient mice compared to gp33 10^-6^ M derived P14 cells, presumably due to the diminished abundance of CD62L^+^TCF1^+^ cells within the transferred bulk population (Fig. 4G). Accordingly, absolute numbers of IL7R^+^KLRG1^-^ P14 memory cells were significantly higher in recipient mice that received C6 or gp33 10^-11^ M stimulated cells (Fig. 4G). P14 cells derived from both weak TCR stimulation conditions were mainly either CD62L^+^TCF1^+^ or CD62L^-^TCF1^+^, whereas P14 cells derived from the strong TCR stimulation condition had similar numbers of CD62L^+^TCF1^+^, CD62L^-^TCF1^+^ and CD62L^-^TCF1^-^ cells (Fig. 4G).

**Figure 4:**
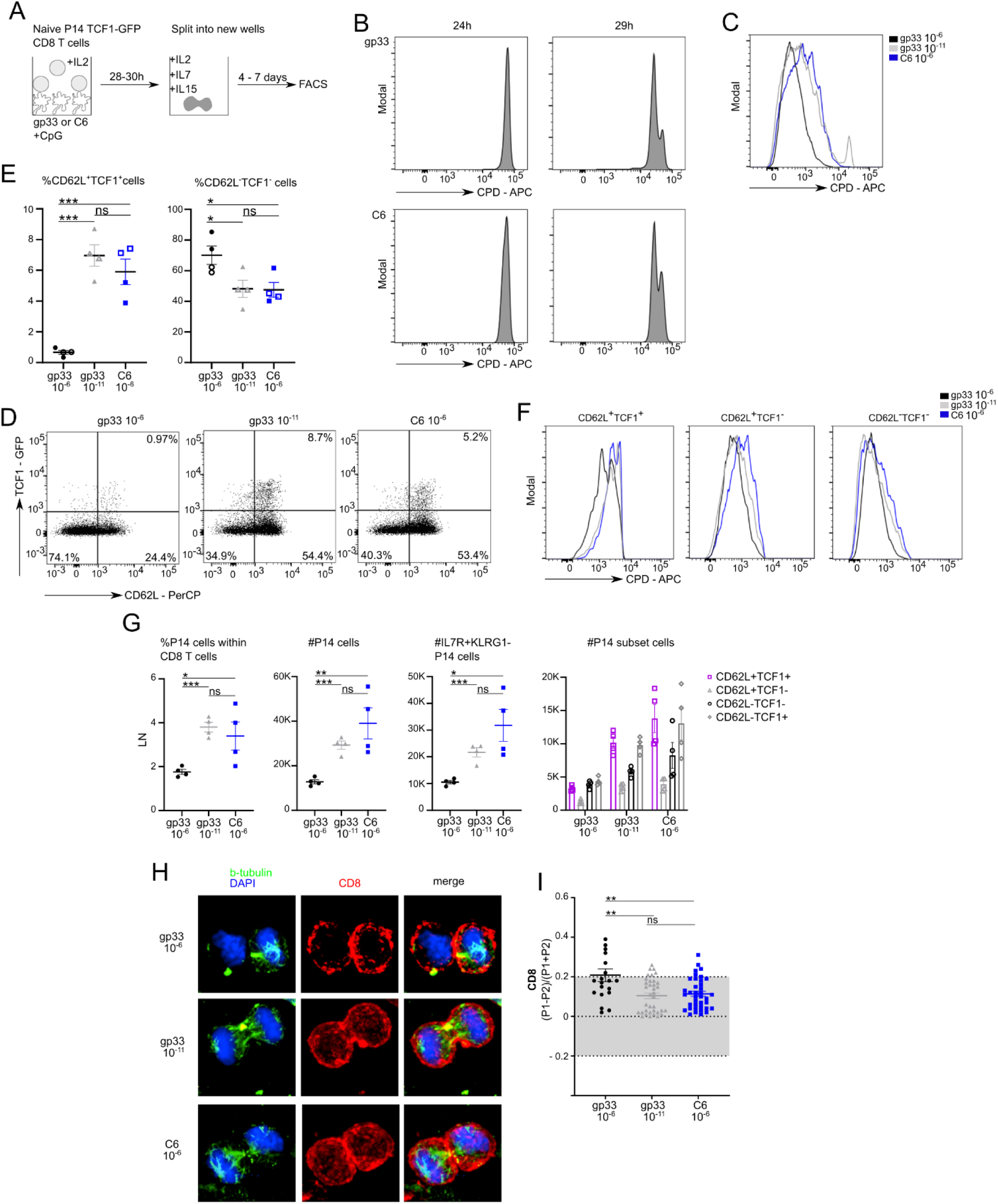
The strength of TCR stimulation impacts ACD and fate. **A** Experimental setup. MutuDC1940 cells were stimulated with CpG and pulsed with either gp33 (high affinity) or C6 (low affinity) peptides before P14 TCF1-GFP cells were added. P14 cells were activated in the presence of IL-2 for 28 – 30 h before cells were either analyzed for ACD or transferred to new wells in medium containing IL-2, IL-7 and IL-15. Cells were cultured for 4 - 7 days. B Representative histograms of CPD dilution of gp33 or C6 activated P14 TCF1-GFP cells after 24 and 29 h. C Representative histograms of CPD dilution of gp33 at 10^-6^ M, gp33 at 10^-11^ M or C6 at 10^-6^ M activated P14 TCF1-GFP cells on day 6. D Representative FACS plots of TCF1 and CD62L expression. P14 TCF1-GFP cells were analyzed on day 6 after activation. E Frequencies of CD62L^+^TCF1^+^ and CD62L^-^TCF1^-^ cells on day 4 (empty symbol) or day 6 (filled symbol) after stimulation. F Histograms of CPD dilution of CD62L^+^TCF1^+^, CD62L^+^TCF1^-^ and CD62L^-^TCF1^-^ cells on day 6 post activation. G P14 TCF1-GFP cells were activated with gp33 at 10^-6^ M, 10^-11^ M or C6 at 10^-6^ M and sorted on day 6 post activation. Sorted cells were individually transferred at equal numbers into recipient mice followed by acute LCMV WE infection (200 ffu/mouse intravenously) one day later. Lymph nodes were harvested 35 days post infection. Frequencies and absolute numbers of P14 cells within lymph nodes of recipient mice. Absolute numbers of IL7R^+^KLRG1^-^, CD62L^+^TCF1^+^, CD62L^+^TCF1^-^, CD62L^-^TCF1^-^ and CD62L^-^TCF1^+^ cells. H Confocal images from fixed samples of naïve P14 cells 27 – 30 h after *in vitro* stimulation with gp33 or C6 loaded MutuDCs at 10^-6^ M or 10^-11^ M. Mitotic cells were identified based on β-tubulin and nuclear structures and imaged from late anaphase to cytokinesis. I ACD rates from P14 cells activated with gp33 at 10^-6^ M (n=20), gp33 at 10^-11^ M (n=36) or C6 at 10^-6^ M (n=38). Data are represented as mean ± SEM. (B) Representative data from one of two experiments. (C, D and F) Representative data from one of three experiments. (E) Data pooled from three independent experiments. (G) Representative data from one of two experiments. Statistical analysis was performed using the unpaired two-tailed Student’s *t* test or, when data did not pass the Shapiro-Wilk normality test, the unpaired two-tailed Mann-Whitney test. \**P* < 0.05; \*\**P* < 0.01; \*\*\**P* < 0.001.

We next addressed the frequency of ACDs after activation by pMHC/DC with different affinities and concentrations. In line with previous reports [11], we found that activation with high concentrations of the high affinity gp33 (10^-6^ M) peptide induced increased frequencies of ACD compared to activation with the C6 variant or gp33 at 10^-11^ M (Fig. 4H and I). To exclude a P14-specific effect, we additionally analyzed first mitosis ACD rates and differentiation after stimulation of OT-I TCF1-GFP cells with their respective high affinity (SIINFEKL “N4”) and low affinity (SIILFEKL “L4”) peptide variants [47]. Also here, stimulation with N4-loaded DCs led to higher frequencies of ACDs compared to stimulation with L4-loaded DCs (Fig. S3D). Additionally, the frequency of CD62L^+^TCF1^+^ cells was lower after N4-stimulation compared to L4-stimulation (Fig. S3E and S3F). Taken together, strong TCR stimulation resulted in increased ACD rates with a preferential path of differentiation into an effector fate. Weak TCR stimulation, in contrast, either provided by low affinity peptides or low antigen concentration, resulted in lower ACD rates and in uniform differentiation into memory or effector precursor cells.

### *In vitro* generated CD62L^+^TCF1^+^ and CD62L^-^TCF1^-^ cells are transcriptionally similar to *in vivo* generated memory and effector cells

Following the observation that strong TCR stimulation leads to enhanced ACD rates and preferential generation of CD62L^-^TCF1^-^ effector precursor cells at the expense of CD62L^+^TCF1^+^ memory precursor cells, which results in curtailed memory formation, we next aimed to dissect the role of ACD on fate diversification specifically upon strong TCR stimulation. To complement the *in vitro* and *in vivo* characterization of CD62L^+^TCF1^+^ and CD62L^-^TCF1^-^ cells derived from different strengths of TCR stimulation on a transcriptional level, we analyzed the transcriptional profiles of *in vitro* generated CD62L^+^TCF1^+^ and CD62L^-^TCF1^-^ cells derived from either antibody-induced (AB) activation or from gp33/C6 peptide stimulation and compared them to *in vivo* generated effector and memory cells, respectively. Thus, we sorted CD62L^+^TCF1^+^ and CD62L^-^TCF1^-^ cells on day 6 post activation, followed by RNA extraction and bulk RNA sequencing. Using multidimensional scaling, we found that biological replicates from each condition clustered closely together and that CD62L^+^TCF1^+^ and CD62L^-^TCF1^-^ samples occupied distinct areas, indicating differential transcriptional profiles. Interestingly, independent of their initial activation stimulus, samples within each subset showed overall transcriptional similarity (Fig. 5A and S4A). Next, we analyzed gene expression of known memory- and effector-specific genes. As expected, *Sell* (encoding CD62L) and *Tcf7* (encoding TCF1) were expressed at higher levels in CD62L^+^TCF1^+^ cells (Fig. 5B and S4B). Effector genes, such as *Gzma*, *Gzmb* and *Tnf* were found to be enriched in CD62L^-^TCF1^-^ cells in all activation conditions (Fig. 5B and S4B). Interestingly, while expression patterns of *Tbx21* and *Ezh2* were similar between AB and C6 activation conditions with higher expression in CD62L^-^TCF1^-^ cells, gp33 activation showed an altered expression pattern with higher or similar expression in CD62L^+^TCF1^+^ cells. Furthermore, genes related to terminal differentiation and inhibitory regulation (e.g., *Pdcd1*, *Ctla4* and *Prdm1*) were predominantly expressed in cells derived from strong gp33 activation, with CD62L^-^TCF1^-^ cells displaying higher expression (Fig. 5B and S4B). Differential gene expression (DEG) analysis between CD62L^+^TCF1^+^ and CD62L^-^TCF1^-^ cells further confirmed transcriptional differences between CD62L^+^TCF1^+^ and CD62L^-^TCF1^-^ cells (Fig. 5C). To investigate whether the DEG relate to *in vivo* generated effector and memory cell transcriptional profiles, we performed gene set enrichment analysis (GSEA) using gene sets from effector CD8 T cells 8 days post infection (dpi) with acute LCMV Armstrong and >40 dpi memory cells, respectively [48]. We found a significant overlap of DEG of CD62L^+^TCF1^+^ cells (positive fold change (FC)) with downregulated DEG of 8 dpi effector cells versus >40 dpi memory cells in all activation conditions (Fig. 5D). Accordingly, we observed a significant enrichment of DEG of CD62L^-^TCF1^-^ cells (negative FC) in upregulated DEG of 8 dpi effector cells versus >40 dpi memory cells in all activation conditions (Fig. 5D). To confirm our findings, we performed another GSEA with gene sets from IL7R^lo^ short-lived effector cells (SLECs) and IL7R^hi^ memory precursor effector cells (MPECs) 6/7 days post LCMV infection, respectively. Similarly, we found a significant overlap of DEG of CD62L^+^TCF1^+^ cells (positive (FC)) with downregulated DEG of IL7R^lo^ SLECS versus IL7R^hi^ MPECS in all activation conditions (Fig. S4C). In accordance, we observed a significant enrichment of DEG of CD62L^-^TCF1^-^ cells (negative FC) in upregulated DEG of IL7R^lo^ SLECS versus IL7R^hi^ MPECS in all activation conditions (Fig. S4C) [49]. Thus, these findings confirm that CD62L^+^TCF1^+^ cells transcriptionally resemble *in vivo* generated memory cells, whereas CD62L^-^TCF1^-^ cells are similar to *in vivo* generated effector cells and this is independent of the strength of the initial TCR stimulation. Of note, overall gene expression patterns appeared to be more similar between CD62L^+^TCF1^+^ and CD62L^-^TCF1^-^ cells derived from AB and C6 stimulation compared to the more distinct transcriptional profile of both subsets derived from gp33 activation, once more supporting the notion that AB stimulation can be classified as rather weak compared to high affinity peptide stimulation. Further, as gp33 stimulated cells show a more “effector-oriented” transcriptional profile compared to AB or C6 stimulated cells, it might be possible that a safeguard process might be beneficial, which prevents some cells from adopting a “full effector program”. This seems much less in demand for a low affinity stimulation.

**Figure 5:**
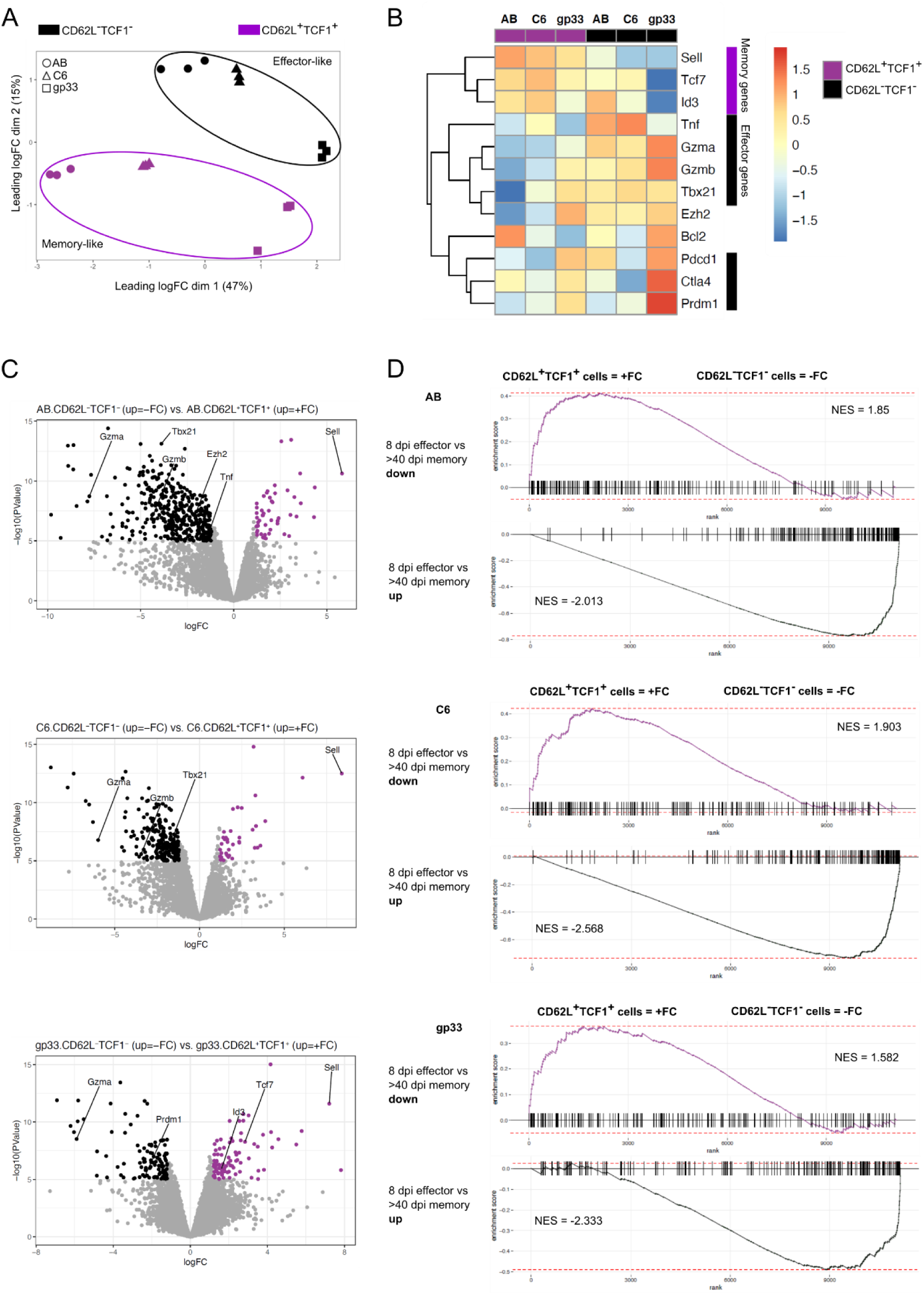
Transcriptional profiling of *in vitro* generated memory and effector precursor cells upon weak and strong TCR stimulation. P14 TCF1-GFP cells were activated by plate-bound Fc-ICAM-1, α-CD3 and α-CD28 (AB) or by MutuDC1940 cells, which were stimulated with CpG and either pulsed with gp33 or C6 peptides. P14 cells were activated in the presence of IL-2 for 30 h before cells were transferred to new wells in medium containing IL-2, IL-7 and IL-15. Cells were cultured for 6 days and CD62L^+^TCF1^+^ and CD62L^-^TCF1^-^ cells were sorted for RNAseq. 3 biological replicates were used for each condition. **A** Multidimensional scaling (MDS) plot depicting sample variation between CD62L^+^TCF1^+^ and CD62L^-^TCF1^-^ cells derived from different stimulation conditions. **B** Heatmap of gene expression of selected memory- and effector-specific genes among CD62L^+^TCF1^+^ and CD62L^-^TCF1^-^ cells derived from AB, C6 or gp33 activation. Averages of pooled biological replicates are shown. **C** Volcano plots displaying differentially expressed genes between CD62L^+^TCF1^+^ and CD62L^-^TCF1^-^ cells per activation condition. Points colored in purple or black are differentially expressed (p-value < 10^-6^ and average log fold change (FC) > 1.2). **D** Gene set enrichment analysis (GSEA) plots and normalized enrichment score (NES) of differential gene expression displayed in C using gene sets describing 8 days post acute infection (dpi) with LCMV Armstrong effector cells and >40 dpi memory cells, respectively (Kaech et al., 2002).

### ACD enables the establishment of single cell-derived mixed-fate colonies upon strong TCR stimulation

To investigate the role of ACD in fate divergence specifically upon strong TCR stimulation at high resolution, we next addressed on a single-cell level whether ACD gives rise to differential cell fates within a single cell-derived colony in this setting. To this end, we activated P14 TCF1-GFP cells by gp33- or C6-loaded DCs for 24 h before we sorted single, undivided but blasted cells into wells of a 384 well plate (sorting strategy shown in Fig. S5A). We then performed a high-throughput microscopy approach using time-lapse imaging with an interval of 60 minutes (min) for the following 24 h to record the first cell division (Movie 3). Cells were then incubated for another 3 days before formed colonies were imaged and analyzed for fate acquisition (Fig. 6A). Quantifying the CD8 surface expression within the first 24 h of imaging demonstrated that cells divided both asymmetrically and symmetrically (Fig. 6B). Strikingly, upon strong TCR stimulation (gp33), we observed that some single cell-derived colonies consisted of both TCF1^+^ memory and TCF1^-^ effector precursor cells on day 5 after stimulation (Fig. 6C and S5C). To exclude that the detected green signal was the result of potential auto-fluorescent debris of dead cells, we performed a propidium iodide (PI) staining, which revealed that the detected signal was not associated with dead cells (Fig. S5B). We next analyzed whether mixed fate colonies were preferentially derived from mother cells undergoing ACD in their first mitosis. Compared to control day 5 sorted and subsequently imaged TCF1^+^ memory precursor cells (green dots) and TCF1^-^ effector precursor cells (brown dots), we observed a significant enrichment of TCF1-GFP^+^ memory precursor cells in colonies stemming from an initial ACD (red dots) compared to colonies emerging from an initial symmetric cell division (SCD) (black dots) (Fig. 6C). However, colonies stemming from a SCD also contained sometimes very few TCF1-GFP^+^ cells (Fig. 6C-E). When comparing the ratio of TCF1-GFP^+^ cells per colony (defined as cells with a GFP signal above the mean value of control sorted TCF1-GFP^+^ cells) to total cell number of the colony, a significant difference was observed between ACD- and SCD-derived colonies with ACD-derived colonies containing more TCF1-GFP^+^ memory precursor cells (Fig. 6D). Consequently, when comparing the ratio of TCF1-GFP^-^ cells per colony (defined as cells with a GFP signal below the mean value of control sorted TCF1-GFP^-^ cells) to total cell number of the colony, SCD-derived colonies contained more TCF1-GFP^-^ effector precursor cells compared to ACD-derived colonies (Fig. 6D). This approach, however, is impacted by the fact that effector precursors proliferate faster than memory precursor cells and therefore influence the absolute cell number per colony. This might lower the ratio of TCF1-GFP^+^ cells and increase the ratio of TCF1-GFP^-^ cells within a colony. We therefore compared the absolute numbers of TCF1-GFP^+^ or TCF1-GFP^-^ cells per colony and found that ACD-derived colonies indeed contained significantly more TCF1-GFP^+^ memory precursor cells compared to SCD-derived colonies (Fig. 6E). SCD-derived colonies, concomitantly, contained significantly more TCF1-GFP^-^ effector cells compared to ACD-derived colonies (Fig. 6E). In addition, we found that single cells, which established mixed-fate colonies, underwent more ACDs during their first cell division compared to those that established single-fate colonies (Fig. 6F). To strengthen our hypothesis and confirm the findings from the plate-bound antibody induced stimulation conditions (Fig. 3, representing a weak TCR stimulation), we repeated the experiment using C6 stimulation. Indeed, single cell-derived colonies preferentially induced either small colonies comprising TCF1-GFP^+^ memory precursor cells or large colonies consisting of TCF1-GFP^-^ effector precursor cells and no mixed-fate colonies (Fig. 6G and S5D). As for the antibody-induced stimulation condition (Fig. 3E and F), we did not observe any correlation between first mitosis ACD or SCD with subsequent fate diversification (Fig. 6H-J). Taken together, our data suggest that ACD serves as a mechanism for the generation of memory precursor cells, specifically upon strong TCR stimulation.

**Figure 6:**
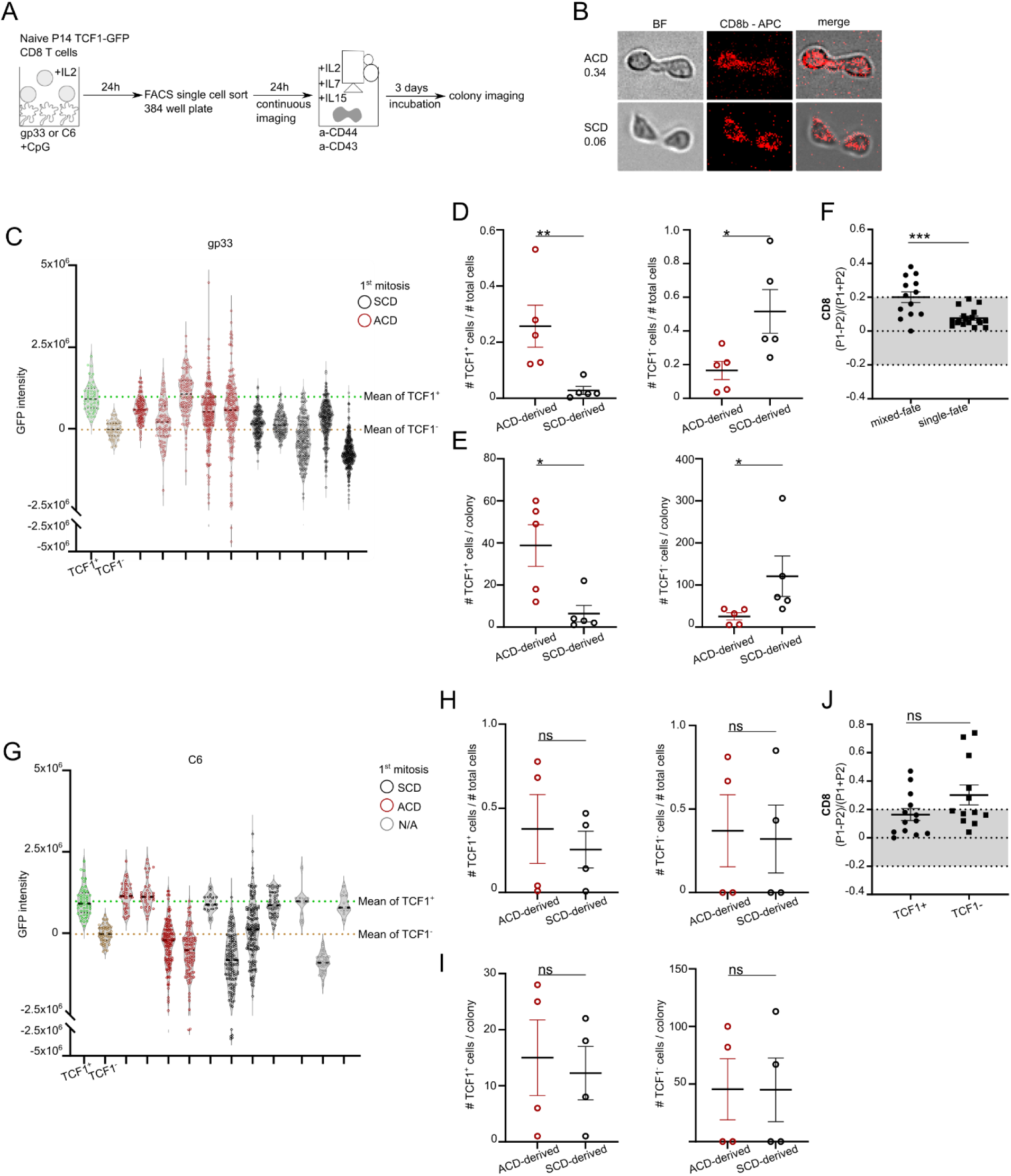
ACD enables the establishment of single cell-derived mixed-fate colonies upon strong TCR stimulation. **A** Experimental setup. MutuDC1940 cells were stimulated with CpG and pulsed with gp33 or C6 peptide before P14 TCF1-GFP cells were added. P14 cells were activated in the presence of IL-2 for 24 h before blasted single cells were sorted into a 384-well plate containing medium supplemented with IL-2, IL-7 and IL-15. Cells were imaged in BF and red channel (CD8b-APC) every 60 min for 24 h and then further incubated. On day 5 post activation established colonies were imaged and analyzed for TCF1 expression. **B** Representative images of P14 TCF1-GFP cells in BF and red channel (CD8b-APC) taken from the time-lapse movie at the first time-point after cell division. **(C-F)** Data derived from gp33 stimulation. **(G-J)** Data derived from C6 stimulation. C+G GFP intensities of single cells within established colonies stemming from an initial ACD (red) or SCD (black). N/A indicates that no reliable asymmetry determination was possible. Green dots: control sorted and imaged TCF1^+^ cells from day 5 post *in vitro* activation. Brown dots: control sorted and imaged TCF1^-^ cells from day 5 post *in vitro* activation. **D+H** Ratio of cell numbers of TCF1^+^ and TCF1^-^ cells divided by total cell numbers per colony. **E+I** Absolute cell numbers of TCF1^+^ and TCF1^-^ cells per colony. F ACD rates from P14 cells activated with gp33 resulting in mixed-fate (n=13) or single-fate (n=18) colonies. A colony was characterized as mixed-fate when more than 10 TCF1^+^ cells were identified. J ACD rates from P14 cells activated with C6 resulting in single-fate TCF1^+^ (n=13) or TCF1^-^ (n=12) colonies. Statistical analysis was performed using the unpaired two-tailed Student’s *t* test or, when data did not pass the Shapiro-Wilk normality test, the unpaired two-tailed Mann-Whitney test. \**P* < 0.05, \*\**P* < 0.01, \*\*\**P* < 0.001.

### Inhibition of ACD markedly curtails memory precursor formation upon strong TCR stimulation

Following the hypothesis that ACD is of particular importance in memory cell generation upon strong TCR stimulation, we postulated that ACD might function as a safeguard mechanism for memory cell generation, specifically upon strong TCR stimulation. To test this assumption, we set out to modulate the ability of cells to undergo ACD in order to investigate the effect on memory formation. We hypothesized that interfering with the ability of cells to perform ACD would not influence fate outcome in conditions of weak TCR stimulation, whereas it would have a clear impact in conditions of strong TCR stimulation. Previous studies have demonstrated that atypical PKC is involved in the regulation of ACD and that PKCζ inhibition impairs the establishment of asymmetry [14], [18], [19], [22]. To this end, we used a myristolated PKCζ inhibitor (PKCi) to prevent ACD. P14 cells were activated as described in Figure 4A and transiently treated with PKCi for the first 30 h of activation. We observed a significant decrease in ACD rates based on CD8 surface distribution between two daughter cells upon gp33 stimulation in the presence of PKCi compared to gp33 stimulation without PKCi. No effect of PKCi treatment on ACD rates was observed upon C6 stimulation (Fig. 7A and B). Additionally, we treated P14 cells with FTY720, a sphingosine-1-phosphate receptor agonist, previously described to inhibit long chain fatty acid ceramide synthesis, altering lymphocyte trafficking and decreasing ACD rates [22], [50]–[53]. Similar to PKCi treatment, ACD rates were significantly reduced upon FTY720 treatment in the gp33 stimulation condition. No impact of FTY720 on ACD rates in the C6 stimulation condition was observed (Fig. S6A). Next, we investigated the expression profile of TCF1 and CD62L on progenies of activated P14 cells to determine potential effects of ACD inhibition on effector and memory precursor cell development. We discriminated between transient (only for the first 30 h) and permanent (throughout the entire culture period) PKCi treatment of P14 cells upon gp33 and C6 stimulation to inhibit ACD not only during first mitoses but also during all subsequent ones. Transient (t) and permanent (p) PKCi treatment significantly decreased the frequency of CD62L^+^TCF1^+^ cells and increased the frequency of CD62L^-^TCF1^-^ cells upon gp33 stimulation. Interestingly, no difference was observed between transient and permanent PKCi treatment (Fig. 7C and D). Transient PKCi treatment upon C6 stimulation had no effect on the generation of CD62L^+^TCF1^+^ cells and CD62L^-^ TCF1^-^ cells compared to untreated cells. Interestingly, and in contrast to gp33 stimulation, permanent PKCi treatment upon C6 stimulation slightly increased the frequency of memory precursor cells at the expense of effector cells (Fig. 7C and D). However, the observed effects of PKCi treatment were more pronounced in the setting of strong TCR stimulation. To confirm these findings on a functional level, we adoptively transferred P14 cells, either untreated, with transient or permanent PKCi treatment on day 6 after gp33 or C6 stimulation into recipient mice, which were subsequently infected with LCMV WE. In line with the *in vitro* findings, transient and permanent PKCi treatment upon gp33 stimulation significantly decreased the frequencies and absolute numbers of transferred P14 cells in lymph nodes and spleen of recipient mice on day 35 post infection, due to the diminished generation of memory precursor cells (Fig. 7E). Consequently, absolute numbers of IL7R^+^KLRG1^-^ P14 cells were also significantly reduced upon transient and permanent PKCi treatment after gp33 activation (Fig. 7E). Neither transient nor permanent PKCi treatment led to alterations in memory formation upon C6 stimulation (Fig. 7E). Thus, ACD during the first mitosis after activation indeed serves as a safeguard mechanism for the establishment of memory precursor cells in conditions of strong TCR stimulation and prevention of ACD in this setting markedly curtails memory formation *in vitro* and *in vivo*.

**Figure 7:**
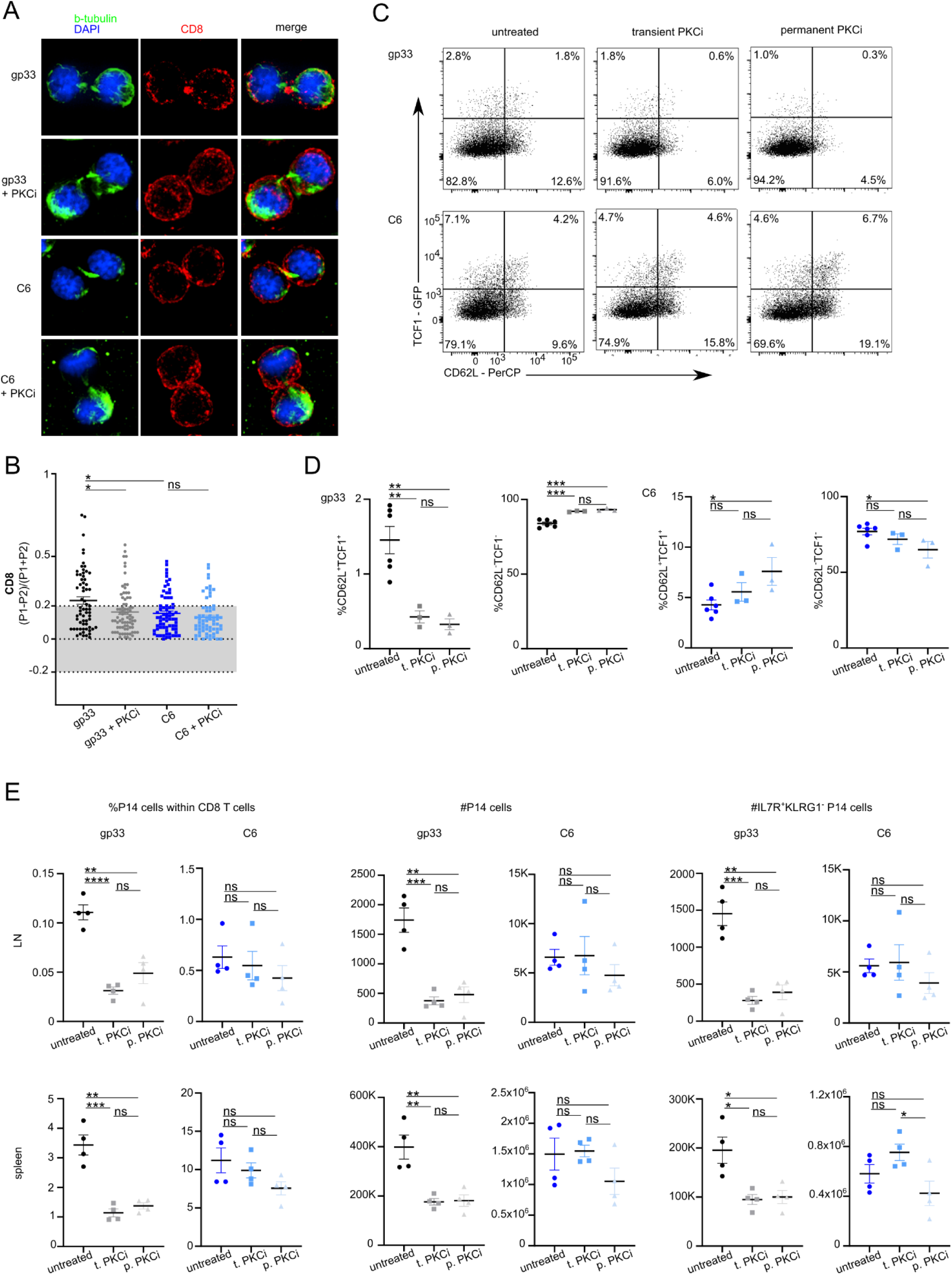
Inhibition of ACD markedly curtails memory precursor formation upon strong TCR stimulation. **A** Confocal images from fixed samples of murine naïve P14 T cells 27 – 30 h after *in vitro* stimulation on gp33 or C6 loaded MutuDCs at 10^-6^ M with or without PKCi treatment. Mitotic cells were identified based on β-tubulin and nuclear structures and imaged from late anaphase to cytokinesis. **B** ACD rates of P14 cells activated with gp33 without (n=65) or with (n=76) PKCi, or with C6 at 10^-6^ M without (n=69) or with (n=65) PKCi. Data are represented as mean ± SEM. **C+D** Representative FACS plots and frequencies of CD62L^+^TCF1^+^ and CD62L^-^TCF1^-^ cells on day 6 post *in vitro* stimulation with gp33 or C6 at 10^-6^ M in the presence or absence of PKCi. PKCi treatment was either transiently (t) performed during the first 30 h of activation or permanently (p) throughout the entire culture period. **E** P14 cells were activated with gp33- or C6-loaded DCs at 10^-6^ M with or without PKCi treatment, either transiently (t) or permanently (p), and sorted on day 7 post activation. Sorted cells were individually transferred at equal numbers into recipient mice followed by acute LCMV WE infection (200 ffu/mouse intravenously) one day later. Lymph nodes and spleens were harvested 35 days post infection. Frequencies and absolute numbers of P14 cells within lymph nodes and spleens of recipient mice. Absolute numbers of IL7R^+^KLRG1^-^ cells. Representative data from 2-3 experiments. Statistical analysis was performed using the unpaired two-tailed Student’s t test or, when data did not pass the Shapiro-Wilk normality test, the unpaired two-tailed Mann-Whitney test. *P < 0.05; **P < 0.01; ***P < 0.001.

## Discussion

The establishment of a heterogeneous pool of CD8 T cell subsets differing in functional, phenotypic and metabolic features is an essential hallmark of effective adaptive immune responses. During acute infections, vigorous proliferation and diversification occur in parallel and it has been reported that a single activated CD8 T cell is able to give rise to both effector and memory cells [6]–[9], [15], [54]. How fate diversification is implemented on a mechanistic level is not fully understood so far. Besides transcriptional and epigenetic regulation, the strength of initial TCR signaling and ACD have gained interest as contributors to the establishment of heterogeneous cell populations. ACD is an evolutionarily well-conserved mechanism for the generation of cellular heterogeneity [55]. In CD8 T cells, ACD is initiated by a stable IS inducing a polarization axis, which allows for unequal distribution of fate-related markers resulting in two daughter cells that differ phenotypically, transcriptionally and metabolically [11], [14], [15], [20]–[22]. Moreover, TCR signaling is mediated via the IS, and modulating the intensity of TCR stimulation has been shown to impact fate diversification, with strong TCR stimulation preferentially inducing effector differentiation and weak TCR stimulation leading to memory precursor formation due to high levels of Eomes and Bcl-6 [11], [12], [56]–[58]. In the present study, we investigated the interplay between ACD and TCR activation signal strength and analyzed their impact on future fate divergence of CD8 T cells on the single cell level. Although previous studies identified remarkable contributions of ACD and TCR signal strength to CD8 T cell differentiation on a bulk population, analysis on the single cell level as well as their interactive contribution was missing. By using different *in vitro* imaging approaches linked with functional *in vivo* analysis and transcriptional profiling, we report a specific safeguarding role for ACD in CD8 memory T cell generation upon strong TCR stimulation. The determination of effector and memory differentiation early after *in vitro* activation requires reliable markers as indicators of fate. In contrast to a recent study describing two separate *in vitro* protocols for the generation of effector and memory precursor cells, respectively, here we established an *in vitro* stimulation protocol which allows for early effector and memory fate discrimination by differential expression of CD62L and TCF1 within the same activation condition [59]. On day 5 post *in vitro* activation, we found that CD62L^+^TCF1^+^ cells show typical hallmarks of memory precursor cells, whereas CD62L^-^TCF1^-^ cells possess features of effector cells. Metabolically, CD62L^+^TCF1^+^ cells relied on oxidative phosphorylation (OXPHOS), whereas CD62L^-^TCF1^-^ cells used glycolysis for energy production. It is well-described that *in vivo* generated memory cells use OXPHOS, while effector cells depend on glycolysis [60]. Consistent with previous studies investigating the proliferation of effector and memory precursor cells, we found that CD62L^+^TCF1^+^ cells divided slower compared to CD62L^-^TCF1^-^ cells [6], [7], [61]–[63]. This finding corresponds to the observed elevated expression levels of CD25 of the CD62L^-^TCF1^-^ cell subset allowing for IL-2 induced proliferation. Furthermore, CD62L^+^TCF1^+^ cells showed nuclear localization of FOXO1, a transcription factor that induces the expression of TCF1 and CD62L and that contributes to orchestrating metabolic regulation favoring the OXPHOS pathway [29]–[31]. By *in vivo* analysis of CD62L^+^TCF1^+^ cells upon adoptive transfer, we found that these cells home better into the T cell zones of lymphoid organs and possess enhanced re-expansion and memory formation potential upon LCMV infection compared to their CD62L^-^TCF1^-^ cell counterpart. Furthermore, the transcriptional profiles of CD62L^+^TCF1^+^ and CD62L^-^TCF1^-^ cells significantly overlapped with *in vivo* established memory and effector cells, respectively. Thus, CD62L^+^TCF1^+^ and CD62L^-^TCF1^-^ cells resemble *in vivo* generated memory and effector cells, respectively, according to phenotypic, transcriptional and functional criteria. Using two LCMV derived peptides with different affinities towards the P14 TCR - gp33 (high affinity) and C6 (low affinity) - allowed us a combined analysis of TCR signal strength and ACD on fate determination [45]. Specifically, our established *in vitro* differentiation protocol enabled us to determine the fate of progenies stemming from a single CD8 T cell a few days after activation - either induced by weak or strong TCR activation - and link the phenotypic composition of the emerging colony to the (a)symmetry of the first cell division of the mother cell. As reported in previous studies, we found that high affinity stimulation induced increased rates of ACD and preferential effector differentiation at the expense of memory formation compared to low affinity stimulation [11], suggesting a potential specific role of ACD within this condition. Therefore, we hypothesized that in contrast to weak TCR stimulation, either induced by low affinity peptides or by plate-bound antibodies, strong TCR stimulation would cause single activated CD8 T cells to generate mixed-fate colonies following ACD. Indeed, we found single cell-derived colonies consisting of effector and memory precursor cells after strong TCR activation. The abundance of TCF1^+^ memory precursor cells was significantly increased if the first performed cell division was asymmetric, while single cell-derived colonies derived from a symmetrically dividing mother cell comprised either none or very few memory precursor cells. In contrast, activation of naïve CD8 T cells by plate-bound Fc-ICAM-1 and α-CD3 and α-CD28 antibodies or by the low affinity C6 peptide did not establish single cell derived mixed-fate colonies. Instead, single cells formed single-fate colonies either consisting of effector (CD62L^-^TCF1^-^) or memory precursor (CD62L^+^TCF1^+^) cells, independent of the (a)symmetry of the first cell division. To confirm a strong TCR stimulation dependent impact of ACD on future fate diversification, we introduced a PKCζ inhibitor into our experimental protocol, leading to inhibition of ACD [14], [18]. We hypothesized that PKCζ inhibition should only impact memory formation after strong TCR stimulation with gp33 and not upon C6 stimulation. Indeed, we found that PKCζ inhibition led to significantly lower ACD rates and strongly curtailed memory formation *in vitro* and *in vivo* after gp33 stimulation, whereas no effect was observed upon C6 stimulation. Lastly, by analyzing permanent versus transient inhibition of ACD by PKCζ inhibitor treatment, we found that ACD during the first mitosis after activation in contrast to potential subsequent ACDs played a key role in fate diversification during CD8 T cell differentiation. Taken together, we report that ACD during the first mitosis after activation functions as a safeguard mechanism for the generation of a robust CD8 memory T cell pool, specifically upon strong TCR stimulation.

Our data and other studies show that strong TCR stimulation induces quantitatively fewer memory precursor cells compared to weak TCR stimulation [11], [12]. It remains to be elucidated, however, whether high-affinity stimulation-derived memory CD8 T cell clones are qualitatively similar or different from low affinity stimulation-derived clones in terms of their responsiveness following recall (i.e. generating potent cytotoxic effector cells and robust secondary memory cells) and are therefore able to compensate for the lower abundance. Interestingly, on a transcriptional level, cells derived from weak TCR stimulation (AB and C6) clustered more closely compared to the more distinct gp33 stimulated cell cluster, which was overall more effector-like. Previous studies provide evidence for differential behavior of low and high affinity memory T cell clones during recall responses, with low affinity memory clones poorly responding to low affinity ligands upon rechallenge [58]. Further, it remains elusive which mechanisms drive ACD upon high versus low affinity stimulation. Even though ACD occurs after weak TCR stimulation at low levels, it does not lead to single cell derived mixed fate colonies, suggesting a differential downstream mechanism that allows fate diversification specifically upon strong TCR stimulation. One option would be the described prolonged interaction between the T cell and the APC, potentially leading to an enforced polarization axis [41]. Whether the recently reported establishment of an ER diffusion barrier forms differentially upon varying TCR signal strengths and thereby potentially contributes to fate divergence remains to be investigated [51]. Furthermore, selective asymmetry of PKCζ could play a fate decisive role as asymmetric PKCζ distribution was exclusively described upon strong TCR stimulation so far [15], [18]. This hypothesis would fit to our results showing that inhibition of PKCζ only affects memory formation after strong TCR stimulation as opposed to weak TCR stimulation. Moreover, it remains to be investigated which mechanisms drive fate divergence upon weak TCR stimulation, as according to our data, ACD does not play a role in this setting. Whether TCR downstream signaling events, such as activation of interleukin-2 inducible tyrosine kinase (ITK), orchestrate a potential threshold of memory versus effector differentiation, which is differentially met by single cells in weak TCR stimulation conditions, but largely overcome in high affinity situations remains to be shown [64]. It should also be noted that the “asymmetry” of a CD8 T cell division is in our study and has been in previous studies at large defined by an arbitrary threshold (often set as a 50% increased distribution of CD8 in one of the two daughter cells), such that functional asymmetry of a CD8 T cell mitosis would be better defined by physiological endpoints. Indeed, our results show that setting this arbitrary threshold would define some cell divisions as being symmetric, despite yielding mixed fate colonies (Fig. 6F). This might be attributed to the fact that asymmetric CD8 inheritance is transient (Fig. S2B) and might therefore have been missed in some of the analyzed mitoses (analysis was only done every 60 min). Thus, a more long-lasting and thus reliable marker of ACD would enable a more precise and long-lasting determination of asymmetric CD8 T cell division and its relevance for future fate determination.

Our imaging approaches allow insights into CD8 T cell differentiation at high resolution. However, emerging new imaging technologies might enable even more precise single-cell tracing experiments, further allowing the combination with downstream manipulations of single cells, such as mitochondrial transfer between living cells, Live-seq or trackSeq [65]–[67]. A general limitation of *in vitro* lineage tracing experiments is the missing link to the physiological *in vivo* environment, which is technically extremely challenging. However, our observation that ACD plays an important role in fate diversification specifically upon strong TCR stimulation provides new perspectives into the regulation of fate diversification during CD8 T cell responses, which can and should in the future be probed and challenged by additional approaches.

## Material and Methods

### Mice

Six to ten-week-old male or female mice were used for the experiments performed in this study. Wild-type Ly5.2 C57BL/6 mice were obtained from the ETH Phenomics Center or from Janvier Labs. C57BL/6J, Ly5.1 P14 mice (CD8^+^ T cells with a transgenic TCR with specificity for the glycoprotein GP_33-41_ epitope of lymphocytic choriomeningitis virus (LCMV) in the context of H-2D^b^ [68], Ly5.1 OTI (CD8^+^ T cells with a transgenic TCR with specificity for the OVA_257-264_ SIINFEKL peptide)) [69] and *Tcf7*^GFP^ reporter mice (expressing GFP under the control of the *Tcf7* locus) [28] were housed and bred under specific pathogen-free conditions in animal facilities at ETH Zurich, Hönggerberg. P14 *Tcf7*^GFP^ and OTI *Tcf7*^GFP^ mice were obtained by crossing *Tcf7*^GFP^ mice to P14 or OTI mice, respectively. All animal experiments were conducted in accordance with the Swiss federal regulations and were approved by the cantonal veterinary office of Zurich (animal experimental permissions: 115/2017, 022/2020).

### Cell lines, virus, viral peptides and infections

Dendritic cells (DC) of the immortalized MuTuDC1940 cell line originate from splenic CD8α conventional DC tumors of C57BL/6J mice and retain all major characteristics of their natural counterparts [46]. LCMV strain WE was kindly provided by R.M. Zinkernagel (University Hospital Zurich), propagated on baby hamster kidney 21 cells [70] and viral titers were determined as previously described [71]. Acute LCMV infections were conducted by injecting 200 ffu of the LCMV WE strain intravenously into the tail vein of recipient mice. LCMV-derived peptides gp33 (KAVYNFATC) and C6 (KAVYNCATC) and OVA-derived peptides N4 (SIINFEKL) and L4 (SIILFEKL) were obtained from EMC Microcollections GmbH.

### CD8 T cell isolation

CD8^+^ T cells were isolated from spleens and lymph nodes of P14 *Tcf7*^GFP^ or OTI *Tcf7*^GFP^ mice using the EasySep^TM^ Mouse CD8+ T cell Isolation Kit (Stemcell Technologies), following manufacturer’s instructions.

### *In vitro* CD8 T cell activation

After isolation, CD8 T cells were kept in complete T cell medium (RPMI-1640 (Bioconcept), 2 mM L-Glutamine (Bioconcept), 2% penicillin-streptomycin (Sigma-Aldrich), 1x Non-Essential Amino Acids (Sigma-Aldrich), 1 mM Sodium Pyruvate (Gibco), 10% fetal bovine serum (Omnilab), 25 mM HEPES (Gibco), 50 µM β-Mercaptoethanol (Gibco)). For antibody induced activation, CD8 T cells were stimulated in presence of self-made human IL-2 on α-CD3 (5 µg/ml) (145-2C11, BioLegend), α-CD28 (5 µg/ml) (37.51, BioLegend) and human Fc-ICAM-1 (50 µg/ml) (R&D Biosciences, Bio-Techne AG) coated plates for 30-36 h at 37°C and 5% CO_2_. To activate CD8 T cells by cognate antigen, adherent MuTuDCs were stimulated with CpG (0.5 µg/ml) and at the same time loaded with the respective peptides at indicated concentrations for 1 h at 37°C and 5% CO_2_. DCs were washed once with PBS before CD8 T cells were added in the presence of self-made human IL-2 and activated for 28-30 h. After initial activation, CD8 T cells were removed from either plate-bound antibodies or adherent dendritic cells and further cultured in complete T cell medium supplemented with IL-7 (10 ng/ml) (eBioscience), IL-15 (5 ng/ml) (eBioscience) and self-made human IL-2. ACD modulation was conducted by adding 30 µM of MYR PKC Zeta Pseudosubstrate (ThermoFisher) to inhibit PKCζ, or by adding 2 µM of FTY720 (Sigma-Aldrich).

### Immunofluorescence staining and confocal microscopy

For tissue section analysis, spleens were incubated in 20% sucrose (Sigma-Aldrich) on a spinning wheel at 4°C for 24 h. Spleens were then embedded in Tissue-Tek® O.C.T. Compound (Sakuraus), snap-frozen in liquid N_2_ and stored at -20°C. 10µm slices were obtained using a microtome. Spleen slices were fixed with 2% paraformaldehyde (PFA) (Sigma-Aldrich) for 1 h at room temperature and blocked with 1×PBS containing 1% of rat serum for 1 h at room temperature. Sections were stained in 1×PBS containing 0.1% of rat serum for 1 h at room temperature in the dark. The following fluorophore-conjugated antibodies were purchased from BioLegend (α-CD8 BV421 53-6.7; α-CD45R/B220 FITC RA3-6B2; α-CD169 (Siglec-1) PE 3D6.112; α-CD45.1 APC A20). Stained slices were mounted on ProLong™ Diamond Antifade Mountant (ThermoFisher) and dried overnight in the dark. For asymmetry analysis, cells were washed with PBS, seeded on poly-l-lysine coated coverslips and incubated for 1 h at 37°C. Fixation of cells was performed by adding 2% PFA (Sigma-Aldrich) for 7 min at room temperature. Cells were then permeabilized with 0.1% Triton X (Sigma-Aldrich) for 10 min and blocked in PBS containing 2% bovine serum albumin (GE Healthcare) and 0.01% Tween 20 (National Diagnostics) for 1 h at room temperature. The following antibodies were used for immunofluorescence staining: mouse α-β-tubulin (Sigma-Aldrich), α-mouse IgG AF488 (Abcam), α-CD8b.2 APC (53-5.8, BioLegend) and α-FOXO1 (C29H4, BioConcept NEB). DAPI (Sigma-Aldrich) was used to identify DNA. Cells were mounted on one drop of ProLong™ Diamond Antifade Mountant (ThermoFisher) onto the microscopy slide and dried overnight in the dark. Microscopic analyses were conducted with a Visitron Confocal System (inverse confocal microscope, Visitron Systems) equipped with four laser lines (405, 488, 561 and 640nm) allowing visualization of four wavelength spectra: blue (405 - 450nm), green (525 - 550nm), red (605 - 652nm) and far-red (700 - 775nm) fluorophores. For each image, 25 Z-stacks (0.5 µm each) were recorded with 10× (spleen sections) (10× objective, type: plan-neofluar, aperture: 0.3, immersion: Air, contrast Ph1) or 100×(asymmetry analysis) magnification (100× objective, type: plan-neofluar, aperture: 1.3, immersion: oil, contrast Ph3) coupled to Evolve 512 EMCCD cameras (Photometrics). Mitotic cells in phases ranging from late anaphase to late telophase were identified based on nuclear morphology, the presence of a dual microtubule-organizing center on each pole of the cell and on characteristic tubulin bridges between two daughter cells in cytokinetic cells (Fig. S8).

### Time-lapse microscopy

For all time-lapse microscopic approaches, cells were cultured at 37°C in a temperature- and CO_2_-controlled incubation chamber in complete T cell medium without phenol red (RPMI-1640 (Gibco), 2 mM L-Glutamine (Bioconcept), 2% penicillin-streptomycin (Sigma-Aldrich), 1x Non-Essential Amino Acids (Sigma-Aldrich), 1 mM Sodium Pyruvate (Gibco), 10% fetal bovine serum (Omnilab), 25 mM HEPES (Gibco), 50 µM β-Mercaptoethanol (Gibco)) supplemented with with IL-7 (10 ng/ml) (eBioscience), IL-15 (5 ng/ml) (eBioscience) and self-made human IL-2. For single-cell continuous imaging after antibody stimulation [34], [35], cells were harvested 32-34 h after activation and transferred into μ-slide VI^0,4^ channel slides (IBIDI), pre-coated with 5 µg/ml α-CD43 (eBioR2/60, Invitrogen) and α-CD44 (IRAWB14) [37]. Movie 1 and 2 were imaged for 3 days using a Nikon-Ti Eclipse equipped with linear-encoded motorized stage, Orca Flash 4.0 V2 (Hamamatsu), and Spectra X fluorescent light source (Lumencor) with CFP (436/20; 455LP; 480/40), GFP (470/40; 495LP; 525/50) and Cy5 (620/60; 660LP; 700/75; all, AHF) filter cubes to detect CD62L-BV421, TCF1-GFP and CD8-APC, respectively. Images were acquired every 40 min in BF and the Cy5 channel and every 90 min in the CFP and GFP channels with a 10× (NA 0.45) CFI Plan Apochromat λ objective. For single-cell high-throughput imaging, cells were harvested 24 h after initial activation and stained for viability and CD8b-APC before blasted single cells were sorted into wells of a 384-well plate pre-coated with 5 µg/ml α-CD43 (eBioR2/60, Invitrogen) and α-CD44 (IRAWB14) [37]. Cells were time-lapse imaged using an ImageXpress Micro Confocal Microscope (Molecular Devices) for 24 h in an interval of 1 h to record the first cell division. After 24 h, cells were removed from the microscope and incubated for another 3 days. Single-cell derived colonies were subsequently imaged for fate analysis. Bright-field and fluorescence images were acquired using the 10× objective using MetaXpress software (Molecular Devices). Microscopy images were analyzed using the ImageJ softwares. Single cell tracking was performed using the tTt software [36].

### Quantification

Wide field and confocal microscopy images were analyzed using the ImageJ software. GFP intensities were measured by subtracting the background signal next to the cell from each individual cell from the signal obtained by the region of interest (ROI) drawn around the cell. Asymmetry rates were calculated based on CD8 signal quantification of both daughter cells (detailed information of determining the ROI in Fig. S7). The raw integrated density (RID) of each daughter cell was first normalized to the respective area. Next, a threshold at 40% was applied on both daughter cells, excluding 60% of dim, potentially unspecific signal [72]. Using this threshold, a new measurement was performed creating updated raw integrated density values. The final formula led us to determine the asymmetry rate (AR) of two daughter cells (Equation 1). Cell divisions were defined as asymmetric when the CD8 signal was 1.5-fold greater in one daughter cell compared to the other, equaling an AR of 0.2.

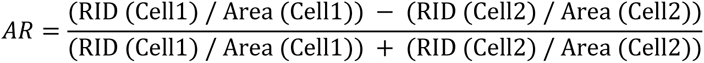

### Flow cytometry

Inguinal lymph nodes, spleens and lungs were obtained from PBS-perfused mice. Spleens and inguinal lymph nodes were smashed through 70 µm strainers (BD Biosciences) using a syringe plunger in order to prepare single-cell suspensions. Lungs were cut into smaller pieces and further incubated in RPMI-1640 (BioConcept) containing 2 mM L-Glutamine (Bioconcept), 2% penicillin-streptomycin (Sigma-Aldrich), 1x Non-Essential Amino Acids (Sigma-Aldrich), 1 mM Sodium Pyruvate (Gibco), 10% fetal bovine serum (Omnilab), 25 mM HEPES (Gibco), 50 µM β-Mercaptoethanol (Gibco) and 2.4 mg/ml collagenase type I (Gibco) and 0.2 mg/ml DNase I (Roche Diagnostics) for 40 min at 37°C. Next, mononuclear cells were obtained by gradient centrifugation over 30% Percoll (Sigma-Aldrich). Erythrocytes were removed by ACK lysis buffer treatment at room temperature for 5 min. When cytokine production was investigated, CD8 T cells were stimulated with 1 µg/ml of gp33 peptide in the presence of 10 µg/ml Brefeldin A (Sigma-Aldrich) at 37°C for 6 h. Fluorophore-conjugated antibodies used for flow cytometry stainings were purchased from BD Biosciences (α-CD278 (ICOS) PE 7E.17G9), eBiosciences (α-TNF Pe-Cy7 TN3-19.12), Miltenyi Biotec (α-phospho-Akt (pS473) PE REA359) and BioLegend (α-CD8b.2 APC 53-5.8; α-CD62L PerCP MEL-14; α-CD45.1 APC A20; α-IL-7Rα BV421 A7R34; α-KLRG1 PE-Cy7 2F1; α-CD44 PE IM7; α-CD8 BV510 53-6.7; α-CD11c PE-Cy7 N418; α-CD25 PE 3C7; α-PD-1 PE-Cy7 29F.1A12; α-IL-2 PE JES6-5H4; α-IFNγ Pacific Blue XMG1.2; α-CD366 (Tim-3) PE RMT3-23; α-CD80 PE 16-10A1; α-CD86 APC GL-1; α-CD40 PE-Cy7 3/23; α-CD11c PerCP N418). Identification of viable cells was done by fixable near-IR dead cell staining (Life Technologies). Surface staining was conducted at 4°C for 20 - 30 min. For intracellular cytokine staining, cells were additionally fixed and permeabilized in 2x FACS Lysis Solution (BD Biosciences) with 0.08% Tween 20 (National Diagnostics) at room temperature for 10 min. Intracellular staining was performed at room temperature in the dark for 30 min. In order to investigate phosphoprotein expression, surface staining was followed by fixation with pre-heated 4% PFA (Sigma-Aldrich) at 37°C for 10 min. Cells were then permeabilized with 90% of ice-cold methanol on ice for 30 min and phosphor-staining was conducted at room temperature in the dark for 40 min. Proliferation was analysed by using the Cell Proliferation Dye eFluor™ 670 or Cell Proliferation Dye eFluor® 450 (eBioscience). Following staining, cell suspensions were washed and stored in PBS containing 2% FBS (Omnilab) and 5 mM of EDTA (Sigma-Aldrich) for acquisition. Multiparameter flow cytometry analysis was performed on a FACS Canto^TM^ (BD Biosciences) cell analyzer and fluorescence-activated cell sorting was performed using a BD FACSAria^TM^ (BD Biosciences) cell sorter with FACS Diva software. Data was analyzed using FlowJo software (FlowJo Enterprise, version 10.0.8, BD Biosciences).

### Adoptive transfer

For adoptive transfer, 10 000 or 1000 sorted CD8 T cells were intravenously injected into naïve young CD45.2 C57BL/6 recipient mice.

### Extracellular flux analysis

Sorted CD8 T cells were seeded into poly-l-lysine pre-coated wells of a 96-well plate (XFe96 cell culture microplates, Agilent) and incubated at 37°C for 1 h before the plate was placed into an Extracellular Flux Analyzer XFe96 (Seahorse, Agilent) for measuring the extracellular acidification rate (ECAR) and the oxygen consumption rate (OCR). For ECAR measurement, the following reagents were diluted in pure XF medium and injected into the plate at the indicated timepoints to achieve the final concentrations: 11.6 mM glucose (Sigma-Aldrich), 0.75 μM Oligomycin (ATP-synthase inhibitor) (Adipogen), 100 mM 2-deoxyglucose (2-DG) (Sigma-Aldrich). Non-glycolytic acidification of media (ECAR following injection of 2-DG) was subtracted to calculate glycolysis (ECAR following injection of glucose) and maximal glycolytic capacity (ECAR following injection of Oligomycin). For OCR measurement, the following reagents were diluted in XF medium containing 25 mM glucose (Sigma-Aldrich), 2 mM L-glutamine (BioConcept) and 1 mM Sodium Pyruvate (Gibco), which was adjusted to pH 7.4 and injected into the plate at the indicated timepoints to achieve the final concentrations: 0.75 μM Oligomycin (ATP-synthase inhibitor) (Adipogen), 1 μM FCCP (chemical uncoupler) (Sigma-Aldrich), 1 μM Rotenone and Antimycin A (electron transport chain inhibitors) (Sigma-Aldrich). Non-mitochondrial respiration (OCR values following injection of Rotenone and Antimycin A) was subtracted from overall basal (OCR values prior to injection of Oligomycin), uncoupled (OCR values following injection of Oligomycin) and maximal respiration (OCR values prior injection of Oligomycin) to calculate basal, uncoupled, and maximal mitochondrial respiration, respectively. ATP production was calculated by subtracting uncoupled mitochondrial respiration from basal mitochondrial respiration.

### Bulk RNAseq

CD62L^+^TCF1^+^ and CD62L^-^TCF1^-^ P14 cells were sorted on day 6 after *in vitro* activation with either plate-bound antibodies or peptide-loaded dendritic cells (MutuDC1940 cell line). RNA was extracted using the RNeasy Micro Kit (Qiagen) according to the manufacturer’s protocol and processed for sequencing at the Functional Genomics Center Zürich (FGCZ). mRNA was sequenced on the Illumina platform. For analysis, RNAseq reads were mapped to the mouse reference genome (Ensembl_GRCm38.75) using the R package Rsubread (v2.8.2) with the default parameters [73]. The featureCounts function was then used to assign mapped reads to genomic features using NCBI Entrez IDs [74]. Differential gene expression analysis was performed using the R package edgeR (v3.36.0) [75]. Principal component analysis was performed using the PCAtools package in R (v2.6.0). Heatmaps were created using the pheatmap package (v1.0.12) and general data visualization was performed using ggplot2 (v3.3.6) and ggrepel (v0.9.1). Gene set enrichment analysis was performed using the R package fgsea (v1.20.0), which uses the adaptive multilevel splitting Monte Carlo approach [76]. Gene sets used: KAECH_DAY8_EFF_VS_MEMORY_CD8_TCELL_DN and KAECH_DAY8_EFF_VS_MEMORY_CD8_TCELL_UP [48]; GSE8678_IL7R_LOW_VS_HIGH_EFF_CD8_TCELL_DN and GSE8678_IL7R_LOW_VS_HIGH_EFF_CD8_TCELL_UP [49].

### Statistical analysis

For statistical analysis the unpaired two-tailed Student’s *t* test or, when data did not pass the Shapiro-Wilk normality test, the unpaired two-tailed Mann-Whitney test was performed using GraphPad Prism Software. Statistical significance was determined with \**P* < 0.05; \*\**P* < 0.01; \*\*\**P* < 0.001; \*\*\*\**P* < 0.0001. For animal experiments, sample size varied between 3 to 4 mice per group for each individual experiment.

## Acknowledgements

We thank R. Spörri for scientific discussion and N. Oetiker and F. Wagen for excellent technical assistance. We thank members of the Oxenius, Joller, Sallusto and Latorre groups for helpful discussions.

## Author contributions

F.G. and A.O. designed the experiments; F.G., D. Stark, A.P., A.C., A.W. and M. Balaz performed the experiments; F.G., D. Shlesinger, A.Y. and A.O. analyzed and interpreted the experiments; M. Borsa, N.B. and T.S. provided scientific input; F.G. and A.O. wrote the manuscript.

## References

1. S. M. Kaech and W. Cui, “Transcriptional control of effector and memory CD8+ T cell differentiation,” Nat Rev Immunol, vol. 12, no. 11, pp. 749–761, Nov. 2012, doi: 10.1038/nri3307.

2. J. B. Johnnidis et al., “Inhibitory signaling sustains a distinct early memory CD8 ^+^ T cell precursor that is resistant to DNA damage,” Sci. Immunol., vol. 6, no. 55, p. eabe3702, Jan. 2021, doi: 10.1126/sciimmunol.abe3702.

3. D. Pais Ferreira et al., “Central memory CD8+ T cells derive from stem-like Tcf7hi effector cells in the absence of cytotoxic differentiation,” Immunity, vol. 53, no. 5, pp. 985–1000.e11, Nov. 2020, doi: 10.1016/j.immuni.2020.09.005.

4. S. M. Kaech, J. T. Tan, E. J. Wherry, B. T. Konieczny, C. D. Surh, and R. Ahmed, “Selective expression of the interleukin 7 receptor identifies effector CD8 T cells that give rise to long-lived memory cells.,” Nat Immunol, vol. 4, no. 12, pp. 1191–1198, Dec. 2003, doi: 10.1038/ni1009.

5. H. K. Chung, B. McDonald, and S. M. Kaech, “The architectural design of CD8+ T cell responses in acute and chronic infection: Parallel structures with divergent fates,” Journal of Experimental Medicine, vol. 218, no. 4, p. e20201730, Apr. 2021, doi: 10.1084/jem.20201730.

6. V. R. Buchholz et al., “Disparate Individual Fates Compose Robust CD8 ^+^ T Cell Immunity,” Science, vol. 340, no. 6132, pp. 630–635, May 2013, doi: 10.1126/science.1235454.

7. C. Gerlach et al., “Heterogeneous Differentiation Patterns of Individual CD8 ^+^ T Cells,” Science, vol. 340, no. 6132, pp. 635–639, May 2013, doi: 10.1126/science.1235487.

8. C. Gerlach et al., “One naive T cell, multiple fates in CD8+ T cell differentiation,” Journal of Experimental Medicine, vol. 207, no. 6, pp. 1235–1246, Jun. 2010, doi: 10.1084/jem.20091175.

9. C. Stemberger et al., “A single naive CD8+ T cell precursor can develop into diverse effector and memory subsets.,” Immunity, vol. 27, no. 6, pp. 985–997, Dec. 2007, doi: 10.1016/j.immuni.2007.10.012.

10. Y. Chen, R. Zander, A. Khatun, D. M. Schauder, and W. Cui, “Transcriptional and Epigenetic Regulation of Effector and Memory CD8 T Cell Differentiation,” Front. Immunol., vol. 9, p. 2826, Dec. 2018, doi: 10.3389/fimmu.2018.02826.

11. C. G. King, S. Koehli, B. Hausmann, M. Schmaler, D. Zehn, and E. Palmer, “T Cell Affinity Regulates Asymmetric Division, Effector Cell Differentiation, and Tissue Pathology,” Immunity, vol. 37, no. 4, pp. 709–720, Oct. 2012, doi: 10.1016/j.immuni.2012.06.021.

12. S. Solouki, W. Huang, J. Elmore, C. Limper, F. Huang, and A. August, “TCR Signal Strength and Antigen Affinity Regulate CD8 ^+^ Memory T Cells,” J.I., vol. 205, no. 5, pp. 1217–1227, Sep. 2020, doi: 10.4049/jimmunol.1901167.

13. T. Capece et al., “A novel intracellular pool of LFA-1 is critical for asymmetric CD8+ T cell activation and differentiation,” Journal of Cell Biology, vol. 216, no. 11, pp. 3817–3829, Nov. 2017, doi: 10.1083/jcb.201609072.

14. J. T. Chang et al., “Asymmetric Proteasome Segregation as a Mechanism for Unequal Partitioning of the Transcription Factor T-bet during T Lymphocyte Division,” Immunity, vol. 34, no. 4, pp. 492– 504, Apr. 2011, doi: 10.1016/j.immuni.2011.03.017.

15. J. T. Chang et al., “Asymmetric T Lymphocyte Division in the Initiation of Adaptive Immune Responses,” Science, vol. 315, no. 5819, pp. 1687–1691, Mar. 2007, doi: 10.1126/science.1139393.

16. M. L. Ciocca, B. E. Barnett, J. K. Burkhardt, J. T. Chang, and S. L. Reiner, “Cutting Edge: Asymmetric Memory T Cell Division in Response to Rechallenge,” J.I., vol. 188, no. 9, pp. 4145–4148, May 2012, doi: 10.4049/jimmunol.1200176.

17. S. Liedmann et al., “Localization of a TORC1-eIF4F translation complex during CD8+ T cell activation drives divergent cell fate,” Molecular Cell, vol. 82, no. 13, pp. 2401–2414.e9, Jul. 2022, doi: 10.1016/j.molcel.2022.04.016.

18. P. J. Metz et al., “Regulation of Asymmetric Division and CD8 ^+^ T Lymphocyte Fate Specification by Protein Kinase Cζ and Protein Kinase Cλ/ι,” J.I., vol. 194, no. 5, pp. 2249–2259, Mar. 2015, doi: 10.4049/jimmunol.1401652.

19. J. Oliaro et al., “Asymmetric Cell Division of T Cells upon Antigen Presentation Uses Multiple Conserved Mechanisms,” J.I., vol. 185, no. 1, pp. 367–375, Jul. 2010, doi: 10.4049/jimmunol.0903627.

20. K. N. Pollizzi et al., “Asymmetric inheritance of mTORC1 kinase activity during division dictates CD8+ T cell differentiation,” Nat Immunol, vol. 17, no. 6, pp. 704–711, Jun. 2016, doi: 10.1038/ni.3438.

21. K. C. Verbist et al., “Metabolic maintenance of cell asymmetry following division in activated T lymphocytes,” Nature, vol. 532, no. 7599, pp. 389–393, Apr. 2016, doi: 10.1038/nature17442.

22. M. Borsa et al., “Modulation of asymmetric cell division as a mechanism to boost CD8 ^+^ T cell memory,” Sci. Immunol., vol. 4, no. 34, p. eaav1730, Apr. 2019, doi: 10.1126/sciimmunol.aav1730.

23. A. Guo et al., “cBAF complex components and MYC cooperate early in CD8+ T cell fate,” Nature, vol. 607, no. 7917, pp. 135–141, Jul. 2022, doi: 10.1038/s41586-022-04849-0.

24. B. Kakaradov et al., “Early transcriptional and epigenetic regulation of CD8+ T cell differentiation revealed by single-cell RNA sequencing,” Nat Immunol, vol. 18, no. 4, pp. 422–432, Apr. 2017, doi: 10.1038/ni.3688.

25. D. Loeffler, F. Schneiter, and T. Schroeder, “Pitfalls and requirements in quantifying asymmetric mitotic segregation,” Annals of the New York Academy of Sciences, vol. 1466, no. 1, pp. 73–82, Apr. 2020, doi: 10.1111/nyas.14284.

26. J. T. Chang, E. J. Wherry, and A. W. Goldrath, “Molecular regulation of effector and memory T cell differentiation,” Nature Immunology, vol. 15, no. 12, pp. 1104–1115, Dec. 2014, doi: 10.1038/ni.3031.

27. D.-M. Zhao et al., “Constitutive activation of Wnt signaling favors generation of memory CD8 T cells.,” J Immunol, vol. 184, no. 3, pp. 1191–1199, Feb. 2010, doi: 10.4049/jimmunol.0901199.

28. D. T. Utzschneider et al., “T Cell Factor 1-Expressing Memory-like CD8+ T Cells Sustain the Immune Response to Chronic Viral Infections,” Immunity, vol. 45, no. 2, pp. 415–427, Aug. 2016, doi: 10.1016/j.immuni.2016.07.021.

29. W. C. Adams et al., “Anabolism-Associated Mitochondrial Stasis Driving Lymphocyte Differentiation over Self-Renewal,” Cell Reports, vol. 17, no. 12, pp. 3142–3152, Dec. 2016, doi: 10.1016/j.celrep.2016.11.065.

30. M. V. Kim, W. Ouyang, W. Liao, M. Q. Zhang, and M. O. Li, “The transcription factor Foxo1 controls central-memory CD8+ T cell responses to infection,” Immunity, vol. 39, no. 2, pp. 286–297, Aug. 2013, doi: 10.1016/j.immuni.2013.07.013.

31. G. J. W. van der Windt and E. L. Pearce, “Metabolic switching and fuel choice during T-cell differentiation and memory development,” Immunol Rev, vol. 249, no. 1, pp. 27–42, Sep. 2012, doi: 10.1111/j.1600-065X.2012.01150.x.

32. R. I. K. Geltink, R. L. Kyle, and E. L. Pearce, “Unraveling the Complex Interplay Between T Cell Metabolism and Function,” Annu Rev Immunol, vol. 36, pp. 461–488, Apr. 2018, doi: 10.1146/annurev-immunol-042617-053019.

33. L. Zhang and P. Romero, “Metabolic Control of CD8(+) T Cell Fate Decisions and Antitumor Immunity.,” Trends Mol Med, vol. 24, no. 1, pp. 30–48, Jan. 2018, doi: 10.1016/j.molmed.2017.11.005.

34. D. Loeffler et al., “Asymmetric organelle inheritance predicts human blood stem cell fate.,” Blood, vol. 139, no. 13, pp. 2011–2023, Mar. 2022, doi: 10.1182/blood.2020009778.

35. D. Loeffler et al., “Asymmetric lysosome inheritance predicts activation of haematopoietic stem cells.,” Nature, vol. 573, no. 7774, pp. 426–429, Sep. 2019, doi: 10.1038/s41586-019-1531-6.

36. O. Hilsenbeck et al., “Software tools for single-cell tracking and quantification of cellular and molecular properties,” Nat Biotechnol, vol. 34, no. 7, pp. 703–706, Jul. 2016, doi: 10.1038/nbt.3626.

37. D. Loeffler et al., “Mouse and human HSPC immobilization in liquid culture by CD43- or CD44-antibody coating,” Blood, vol. 131, no. 13, pp. 1425–1429, Mar. 2018, doi: 10.1182/blood-2017-07-794131.

38. M. Borsa et al., “Asymmetric cell division shapes naive and virtual memory T-cell immunity during ageing,” Nat Commun, vol. 12, no. 1, p. 2715, Dec. 2021, doi: 10.1038/s41467-021-22954-y.

39. J. M. Conley, M. P. Gallagher, and L. J. Berg, “T Cells and Gene Regulation: The Switching On and Turning Up of Genes after T Cell Receptor Stimulation in CD8 T Cells,” Frontiers in Immunology, vol. 7, 2016, [Online]. Available: https://www.frontiersin.org/article/10.3389/fimmu.2016.00076

40. A. E. Denton et al., “Affinity thresholds for naive CD8+ CTL activation by peptides and engineered influenza A viruses.,” J Immunol, vol. 187, no. 11, pp. 5733–5744, Dec. 2011, doi: 10.4049/jimmunol.1003937.

41. A. J. Ozga et al., “pMHC affinity controls duration of CD8+ T cell-DC interactions and imprints timing of effector differentiation versus expansion.,” J Exp Med, vol. 213, no. 12, pp. 2811–2829, Nov. 2016, doi: 10.1084/jem.20160206.

42. M. Shakiba et al., “TCR signal strength defines distinct mechanisms of T cell dysfunction and cancer evasion,” Journal of Experimental Medicine, vol. 219, no. 2, p. e20201966, Feb. 2022, doi: 10.1084/jem.20201966.

43. D. Skokos et al., “Peptide-MHC potency governs dynamic interactions between T cells and dendritic cells in lymph nodes,” Nature Immunology, vol. 8, no. 8, pp. 835–844, Aug. 2007, doi: 10.1038/ni1490.

44. J. Zikherman and B. Au-Yeung, “The role of T cell receptor signaling thresholds in guiding T cell fate decisions,” Curr Opin Immunol, vol. 33, pp. 43–48, Apr. 2015, doi: 10.1016/j.coi.2015.01.012.

45. D. T. Utzschneider et al., “High antigen levels induce an exhausted phenotype in a chronic infection without impairing T cell expansion and survival,” J Exp Med, vol. 213, no. 9, pp. 1819–1834, Aug. 2016, doi: 10.1084/jem.20150598.

46. S. A. Fuertes Marraco et al., “Novel murine dendritic cell lines: a powerful auxiliary tool for dendritic cell research.,” Front Immunol, vol. 3, p. 331, 2012, doi: 10.3389/fimmu.2012.00331.

47. M. J. Turner, E. R. Jellison, E. G. Lingenheld, L. Puddington, and L. Lefrançois, “Avidity maturation of memory CD8 T cells is limited by self-antigen expression.,” J Exp Med, vol. 205, no. 8, pp. 1859– 1868, Aug. 2008, doi: 10.1084/jem.20072390.

48. S. M. Kaech, S. Hemby, E. Kersh, and R. Ahmed, “Molecular and Functional Profiling of Memory CD8 T Cell Differentiation,” Cell, vol. 111, no. 6, pp. 837–851, Dec. 2002, doi: 10.1016/S0092-8674(02)01139-X.

49. N. S. Joshi et al., “Inflammation directs memory precursor and short-lived effector CD8(+) T cell fates via the graded expression of T-bet transcription factor.,” Immunity, vol. 27, no. 2, pp. 281– 295, Aug. 2007, doi: 10.1016/j.immuni.2007.07.010.

50. E. V. Berdyshev et al., “FTY720 inhibits ceramide synthases and up-regulates dihydrosphingosine 1-phosphate formation in human lung endothelial cells.,” J Biol Chem, vol. 284, no. 9, pp. 5467–5477, Feb. 2009, doi: 10.1074/jbc.M805186200.

51. H. Emurla, Y. Barral, and A. Oxenius, “Role of mitotic diffusion barriers in regulating the asymmetric division of activated CD8 T cells,” Immunology, preprint, Sep. 2021. doi: 10.1101/2021.09.10.458880.

52. S. Mandala et al., “Alteration of lymphocyte trafficking by sphingosine-1-phosphate receptor agonists.,” Science, vol. 296, no. 5566, pp. 346–349, Apr. 2002, doi: 10.1126/science.1070238.

53. M. Matloubian et al., “Lymphocyte egress from thymus and peripheral lymphoid organs is dependent on S1P receptor 1.,” Nature, vol. 427, no. 6972, pp. 355–360, Jan. 2004, doi: 10.1038/nature02284.

54. C. R. Plumlee, B. S. Sheridan, B. B. Cicek, and L. Lefrançois, “Environmental cues dictate the fate of individual CD8+ T cells responding to infection.,” Immunity, vol. 39, no. 2, pp. 347–356, Aug. 2013, doi: 10.1016/j.immuni.2013.07.014.

55. B. Sunchu and C. Cabernard, “Principles and mechanisms of asymmetric cell division,” Development, vol. 147, no. 13, p. dev167650, Jul. 2020, doi: 10.1242/dev.167650.

56. S. S. Chin et al., “T cell receptor and IL-2 signaling strength control memory CD8+ T cell functional fitness via chromatin remodeling,” Nat Commun, vol. 13, no. 1, p. 2240, Dec. 2022, doi: 10.1038/s41467-022-29718-2.

57. I. Kavazović et al., “Eomes broadens the scope of CD8 T-cell memory by inhibiting apoptosis in cells of low affinity,” PLOS Biology, vol. 18, no. 3, p. e3000648, Mar. 2020, doi: 10.1371/journal.pbio.3000648.

58. K. M. Knudson, N. P. Goplen, C. A. Cunningham, M. A. Daniels, and E. Teixeiro, “Low-Affinity T Cells Are Programmed to Maintain Normal Primary Responses but Are Impaired in Their Recall to Low-Affinity Ligands,” Cell Reports, vol. 4, no. 3, pp. 554–565, Aug. 2013, doi: 10.1016/j.celrep.2013.07.008.

59. V. Neitzke-Montinelli et al., “Differentiation of Memory CD8 T Cells Unravel Gene Expression Pattern Common to Effector and Memory Precursors.,” Front Immunol, vol. 13, p. 840203, 2022, doi: 10.3389/fimmu.2022.840203.

60. E. L. Pearce, M. C. Poffenberger, C.-H. Chang, and R. G. Jones, “Fueling immunity: insights into metabolism and lymphocyte function.,” Science, vol. 342, no. 6155, p. 1242454, Oct. 2013, doi: 10.1126/science.1242454.

61. I. Kinjyo et al., “Real-time tracking of cell cycle progression during CD8+ effector and memory T-cell differentiation,” Nat Commun, vol. 6, no. 1, p. 6301, May 2015, doi: 10.1038/ncomms7301.

62. L. Kretschmer et al., “Differential expansion of T central memory precursor and effector subsets is regulated by division speed,” Nat Commun, vol. 11, no. 1, p. 113, Dec. 2020, doi: 10.1038/s41467-019-13788-w.

63. M. Plambeck et al., “Heritable changes in division speed accompany the diversification of single T cell fate,” Immunology, preprint, Jul. 2021. doi: 10.1101/2021.07.28.454102.

64. J. M. Conley, M. P. Gallagher, A. Rao, and L. J. Berg, “Activation of the Tec Kinase ITK Controls Graded IRF4 Expression in Response to Variations in TCR Signal Strength.,” J Immunol, vol. 205, no. 2, pp. 335–345, Jul. 2020, doi: 10.4049/jimmunol.1900853.

65. W. Chen et al., “Live-seq enables temporal transcriptomic recording of single cells,” Nature, vol. 608, no. 7924, pp. 733–740, Aug. 2022, doi: 10.1038/s41586-022-05046-9.

66. C. G. Gäbelein et al., “Mitochondria transplantation between living cells,” PLOS Biology, vol. 20, no. 3, p. e3001576, Mar. 2022, doi: 10.1371/journal.pbio.3001576.

67. A. Wehling et al., “Combined single-cell tracking and omics improves blood stem cell fate regulator identification,” Blood, Jul. 2022, doi: 10.1182/blood.2022016880.

68. H. Pircher et al., “T cell receptor (TcR) beta chain transgenic mice: studies on allelic exclusion and on the TcR+ gamma/delta population.,” Eur J Immunol, vol. 20, no. 2, pp. 417–424, Feb. 1990, doi: 10.1002/eji.1830200227.

69. K. A. Hogquist, S. C. Jameson, W. R. Heath, J. L. Howard, M. J. Bevan, and F. R. Carbone, “T cell receptor antagonist peptides induce positive selection.,” Cell, vol. 76, no. 1, pp. 17–27, Jan. 1994, doi: 10.1016/0092-8674(94)90169-4.

70. R. Ahmed, A. Salmi, L. D. Butler, J. M. Chiller, and M. B. Oldstone, “Selection of genetic variants of lymphocytic choriomeningitis virus in spleens of persistently infected mice. Role in suppression of cytotoxic T lymphocyte response and viral persistence.,” J Exp Med, vol. 160, no. 2, pp. 521–540, Aug. 1984, doi: 10.1084/jem.160.2.521.

71. M. Battegay, S. Cooper, A. Althage, J. Bänziger, H. Hengartner, and R. M. Zinkernagel, “Quantification of lymphocytic choriomeningitis virus with an immunological focus assay in 24- or 96-well plates.,” J Virol Methods, vol. 33, no. 1–2, pp. 191–198, Jun. 1991, doi: 10.1016/0166-0934(91)90018-u.

72. R. Shimoni, K. Pham, M. Yassin, M. J. Ludford-Menting, M. Gu, and S. M. Russell, “Normalized Polarization Ratios for the Analysis of Cell Polarity,” PLoS ONE, vol. 9, no. 6, p. e99885, Jun. 2014, doi: 10.1371/journal.pone.0099885.

73. Y. Liao, G. K. Smyth, and W. Shi, “The R package Rsubread is easier, faster, cheaper and better for alignment and quantification of RNA sequencing reads.,” Nucleic Acids Res, vol. 47, no. 8, p. e47, May 2019, doi: 10.1093/nar/gkz114.

74. Y. Liao, G. K. Smyth, and W. Shi, “featureCounts: an efficient general purpose program for assigning sequence reads to genomic features.,” Bioinformatics, vol. 30, no. 7, pp. 923–930, Apr. 2014, doi: 10.1093/bioinformatics/btt656.

75. A. T. L. Lun, Y. Chen, and G. K. Smyth, “It’s Delicious: A Recipe for Differential Expression Analyses of RNA-seq Experiments Using Quasi-Likelihood Methods in edgeR.,” Methods Mol Biol, vol. 1418, pp. 391–416, 2016, doi: 10.1007/978-1-4939-3578-9_19.

76. G. Korotkevich, V. Sukhov, N. Budin, B. Shpak, M. N. Artyomov, and A. Sergushichev, “Fast gene set enrichment analysis,” bioRxiv, 2021, doi: 10.1101/060012.

