## Supplemental figures for "Asymmetric cell division safeguards memory CD8 T cell development"

### Supplementary information

### Supplementary Figures

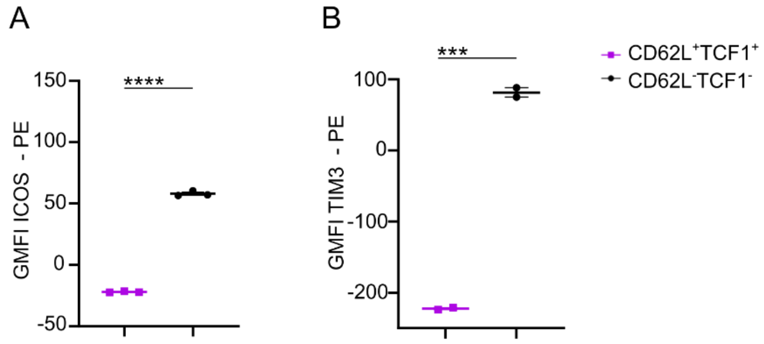


**Figure S1:** ***In vitro*** **differentiation of P14 TCF1-GFP cells and characterization of CD62L$^+$TCF1$^+$ and CD62L$^-$TCF1$^-$ cells.** Geometric Mean of Fluorescence Intensity (GMFI) of CD62L$^+$TCF1$^+$ and CD62L$^-$TCF1$^-$ cells expressing ICOS **(A)** and TIM3 **(B)** on day 4 post stimulation. Statistical analysis was performed using the unpaired two-tailed Student's *t* test or, when data did not pass the Shapiro-Wilk normality test, the unpaired two-tailed Mann-Whitney test. *$P < 0.05$; **$P < 0.01$; ***$P < 0.001$; ****$P < 0.0001$.

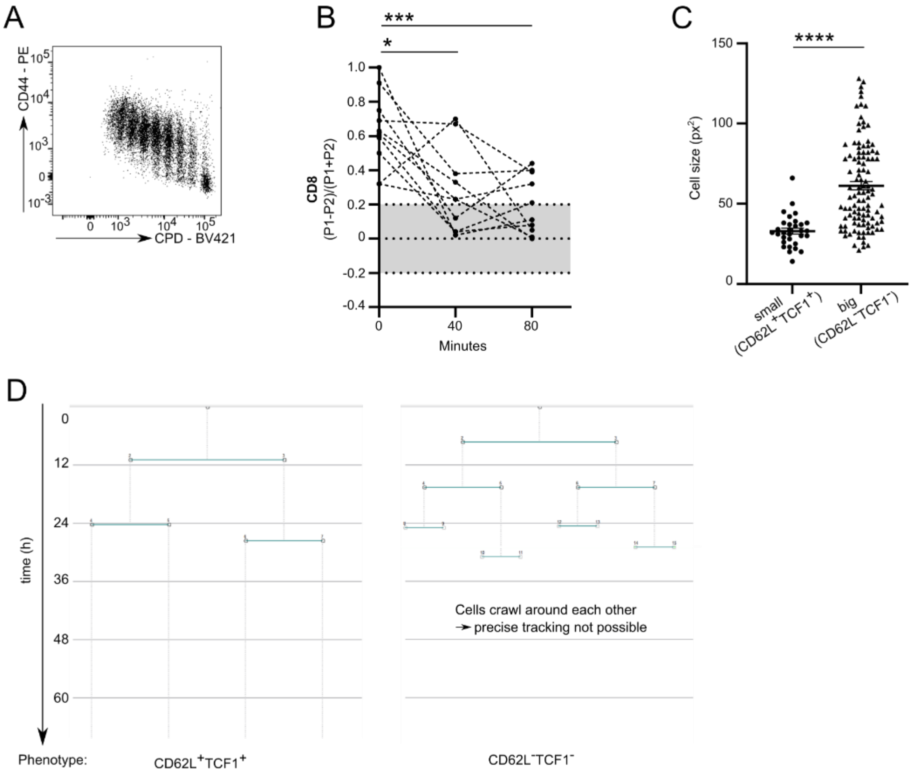


**Figure S2: ACD does not promote diversity upon stimulation with TCR agonistic antibodies.**

**A** Naïve P14 TCF1-GFP cells were stimulated as described in Figure 1A. Representative plot of CPD dilution against expression of CD44 on day 4 post activation. **B** ACD rates of sister cell pairs (n=10) measured directly after completed mitosis, 40 and 80 min later. **C** Cell size (in square pixels) from P14 TCF1-GFP cells that later formed small-sized ($CD62L^{+}TCF1^{+}$) (n=31) and big-sized ($CD62L^{-}TCF1^{-}$) (n=108) colonies. Data are represented as mean ± SEM. **D** Representative family trees of one small-sized colony consisting of $CD62L^{+}TCF1^{+}$ cells and one big-sized colony consisting of $CD62L^{-}TCF1^{-}$ cells. Statistical analysis was performed using the paired (in B) or unpaired (for C) two-tailed Student's *t* test or, when data did not pass the Shapiro-Wilk normality test, the unpaired two-tailed Mann-Whitney test. *$P < 0.05$; **$P < 0.01$; ***$P < 0.001$; ****$P < 0.0001$.

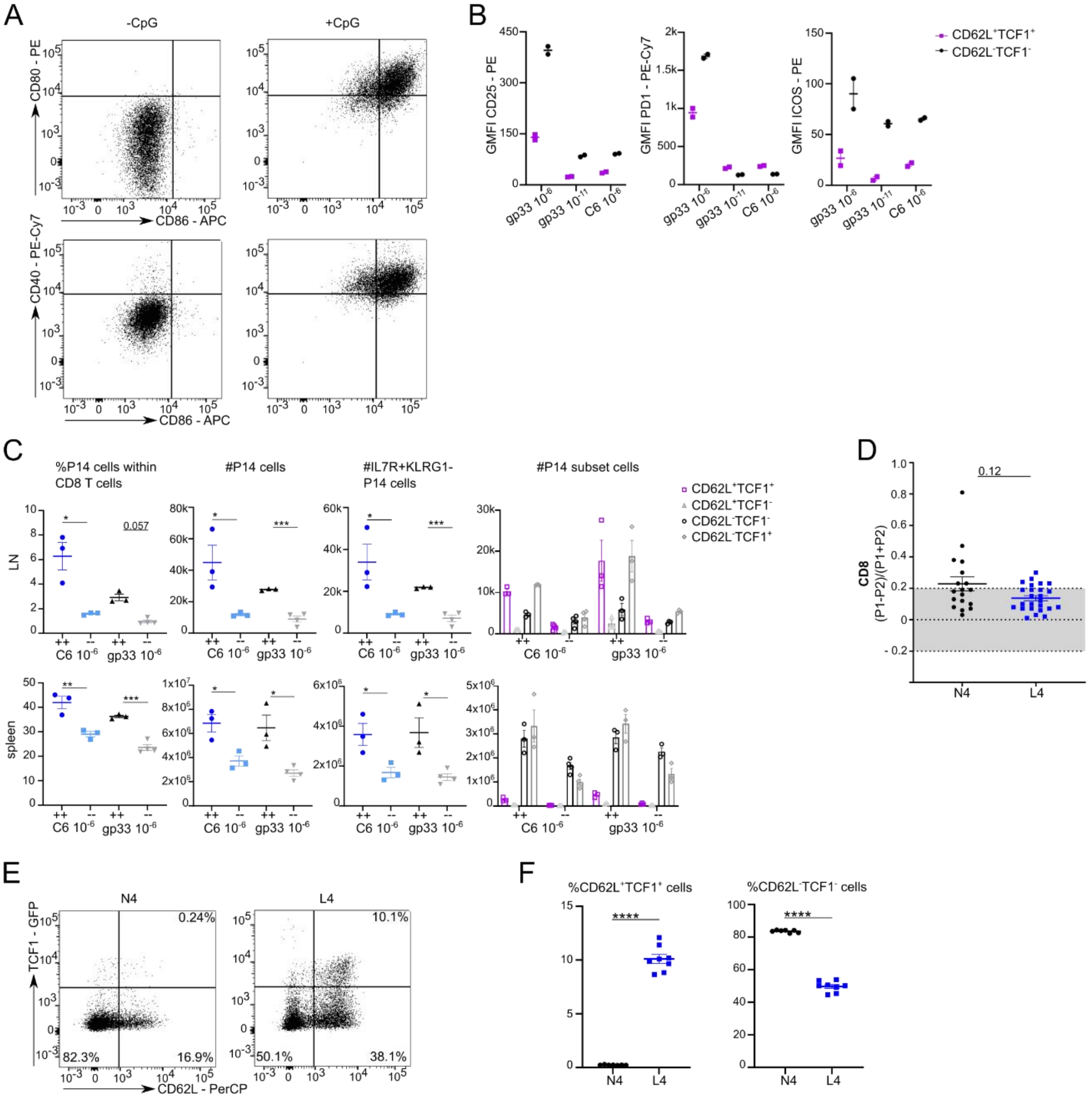


**Figure S3: The strength of TCR stimulation impacts ACD and fate.**

**A** MutuDC1940 cells were either stimulated with or without CpG overnight at 37°C. Representative FACS plots of CD80, CD86 and CD40 expression. **B** Geometric Mean of Fluorescence Intensity (GMFI) of CD25, PD1 and ICOS expression by $CD62L^{+}TCF1^{+}$ and $CD62L^{-}TCF1^{-}$ cells on day 6 post stimulation with gp33 at $10^{-6}$ M, gp33 at $10^{-11}$ M or C6 at $10^{-6}$ M. **C** P14 TCF1-GFP cells were activated with gp33 at $10^{-6}$ M or C6 at $10^{-6}$ M and sorted on day 6 post activation into $CD62L^{+}TCF1^{+}$ and $CD62L^{-}TCF1^{-}$ cells. Sorted subsets were individually transferred at equal numbers into recipient mice followed by acute LCMV WE infection (200 ffu/mouse intravenously) one day later. Spleens, lymph nodes and lungs were harvested 35 days post infection. Frequencies and absolute numbers of P14 cells within spleens and lymph nodes of recipient mice. Absolute numbers of $IL7R^{+}KLRG1^{-}$, $CD62L^{+}TCF1^{+}$, $CD62L^{+}TCF1^{-}$, $CD62L^{-}TCF1^{-}$ and $CD62L^{-}TCF1^{+}$ cells. **D** ACD rates from OT-I cells activated with SIINFEKL (N4) (n=18) or SIILFEKL (L4) (n=25). Data are represented as mean ± SEM. **E** Representative FACS plots of TCF1 and CD62L expression. OT-I TCF1-GFP cells were activated and analyzed on day 6. **F** Frequencies of $CD62L^{+}TCF1^{+}$ and $CD62L^{-}TCF1^{-}$ OT-I cells on day 6 after stimulation. Statistical analysis was performed using the unpaired two-tailed Student's *t* test or, when data did not pass the Shapiro-Wilk normality test, the unpaired two-tailed Mann-Whitney test. *$P < 0.05$; **$P < 0.01$; ***$P < 0.001$.

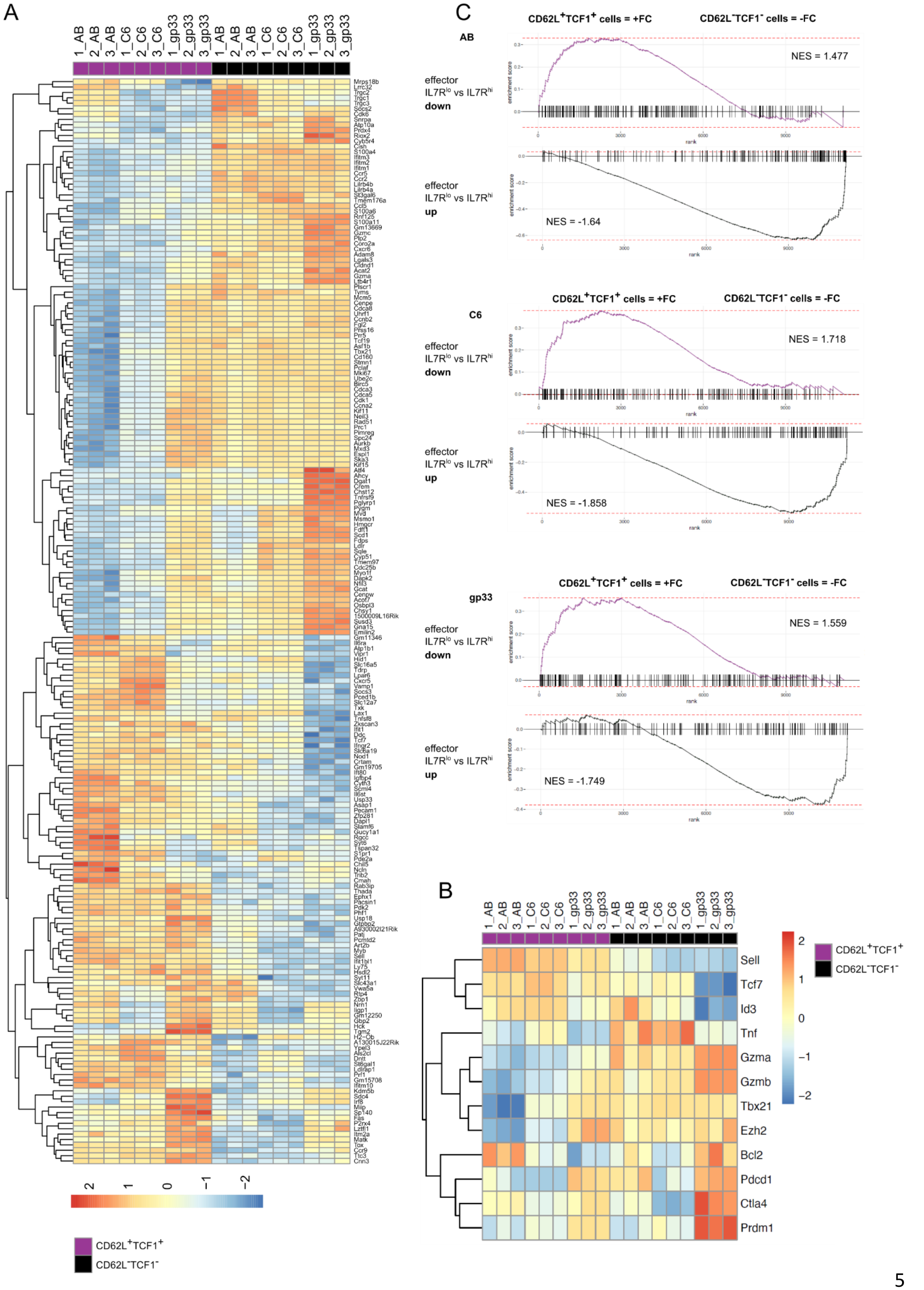
A
1_AB 2_AB 3_AB 1_C6 2_C6 3_C6 1_gp33 2_gp33 3_gp33 1_AB 2_AB 3_AB 1_C6 2_C6 3_C6 1_gp33 2_gp33 3_gp33
2 1 0 −1 −2
CD62L+TCF1+
CD62L−TCF1−
C
CD62L+TCF1+ cells = +FC
CD62L−TCF1− cells = -FC
AB
effector IL7Rlo vs IL7Rhi down
NES = 1.477
effector IL7Rlo vs IL7Rhi up
NES = -1.64
C6
effector IL7Rlo vs IL7Rhi down
NES = 1.718
effector IL7Rlo vs IL7Rhi up
NES = -1.858
gp33
effector IL7Rlo vs IL7Rhi down
NES = 1.559
effector IL7Rlo vs IL7Rhi up
NES = -1.749
enrichment score
rank
B
1_AB 2_AB 3_AB 1_C6 2_C6 3_C6 1_gp33 2_gp33 3_gp33 1_AB 2_AB 3_AB 1_C6 2_C6 3_C6 1_gp33 2_gp33 3_gp33
Sell
Tcf7
Id3
Tnf
Gzma
Gzmb
Tbx21
Ezh2
Bcl2
Pdcd1
Ctla4
Prdm1
2 1 0 −1 −2
CD62L+TCF1+
CD62L−TCF1−

**Figure S4: Transcriptional profiling of *in vitro* generated memory and effector precursor cells upon weak and strong TCR stimulation.**

P14 TCF1-GFP cells were activated by plate-bound Fc-ICAM-1, α-CD3 and α-CD28 (AB) or by MutuDC1940 cells, which were stimulated with CpG and either pulsed with gp33 or C6 peptides. P14 cells were cultured in the presence of IL-2 for 30 h before cells were transferred to new wells in medium containing IL-2, IL-7 and IL-15. Cells were cultured for 6 days and $CD62L^{+}TCF1^{+}$ and $CD62L^{-}TCF1^{-}$ cells were sorted for RNAseq. 3 biological replicates were used for each condition. **A** Differential expression of genes among $CD62L^{+}TCF1^{+}$ and $CD62L^{-}TCF1^{-}$ cells derived from AB, C6 or gp33 activation (top and bottom 50 differentially expressed genes based on the lowest P value) presented as a heatmap. **B** Heatmap of gene expression of selected memory- and effector-specific genes of $CD62L^{+}TCF1^{+}$ and $CD62L^{-}TCF1^{-}$ cells derived from AB, C6 or gp33 activation. Individual biological replicates are shown. **C** Gene set enrichment analysis (GSEA) plots and normalized enrichment score (NES) of differential gene expression displayed in Fig. 5C using gene sets describing 6/7 days post acute LCMV infection $IL7R^{lo}$ short-lived effector cells (SLECs) and $IL7R^{hi}$ memory precursor effector cells (MPECs), respectively (GSE8678) (Joshi et al., 2007).

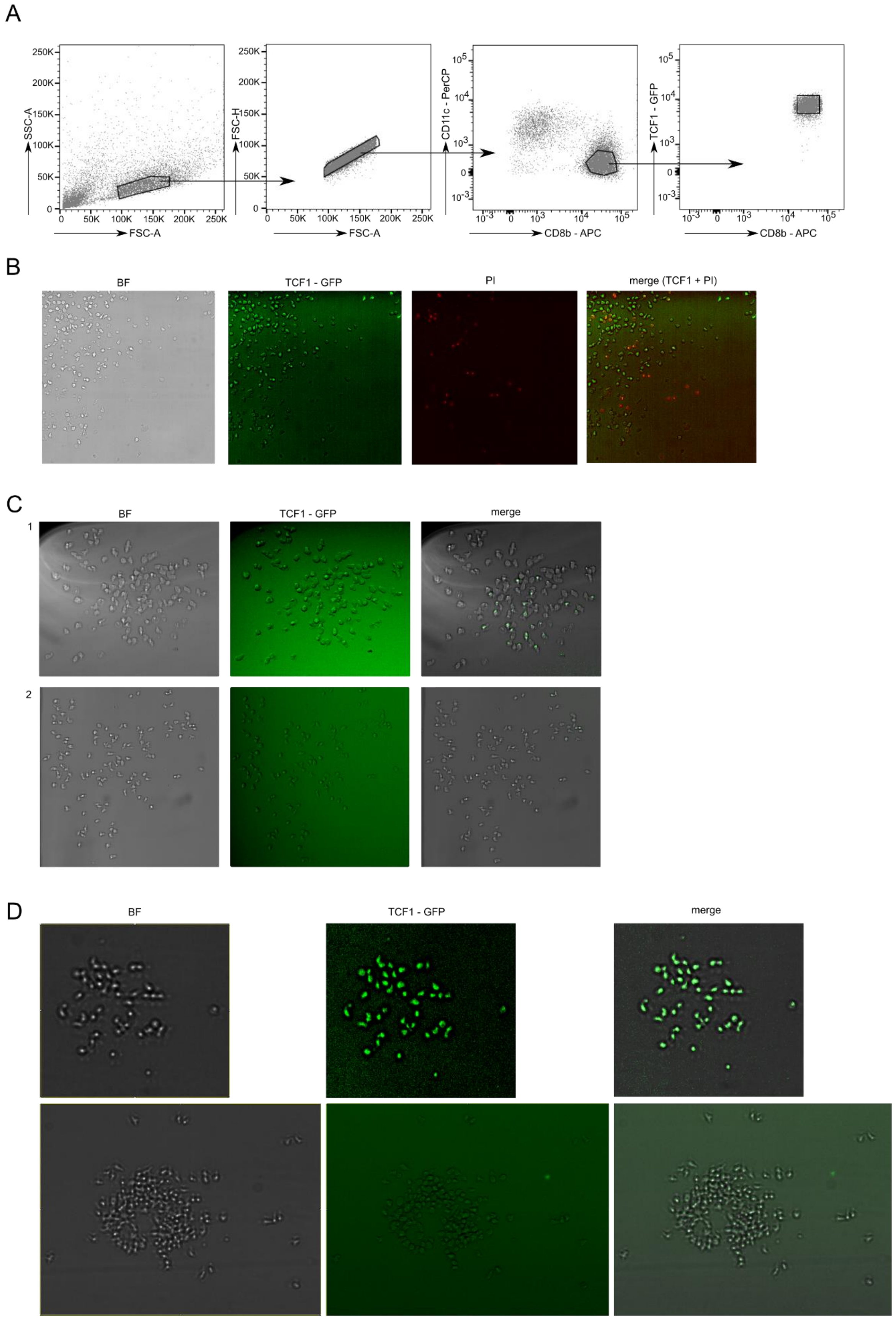
A
SSC-A
FSC-A
FSC-H
FSC-A
CD11c - PerCP
CD8b - APC
TCF1 - GFP
CD8b - APC
B
BF
TCF1 - GFP
PI
merge (TCF1 + PI)
C
BF
TCF1 - GFP
merge
1
2
D
BF
TCF1 - GFP
merge

Figure S5: Strong TCR stimulation results in single cell-derived mixed-fate colonies.

A Gating strategy for sorting of blasted P14 CD8 TCF1-GFP cells 24 h after stimulation. B Propidium Iodide (PI) staining of colonies on day 5 after initial stimulation. C Representative BF and green channel (TCF1-GFP) images of established P14 TCF1-GFP cell colonies stemming from one single mother cell on day 5 post gp33 activation. D Representative BF and green channel (TCF1-GFP) images of established P14 TCF1-GFP cell colonies stemming from one single mother cell on day 5 post C6 activation.

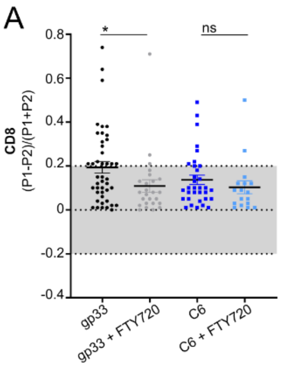


**Figure S6: FTY720 treatment inhibits ACD upon strong TCR stimulation.**

**A** ACD rates from P14 cells activated with gp33 or C6 in the presence or absence of 2 µM FTY720. Data are represented as mean ± SEM (gp33, n=44; gp33 + FTY720, n=25; C6, n=33; C6 + FTY720, n=17). Statistical analysis was performed using the unpaired two-tailed Student's *t* test or, when data did not pass the Shapiro-Wilk normality test, the unpaired two-tailed Mann-Whitney test. *$P < 0.05$.

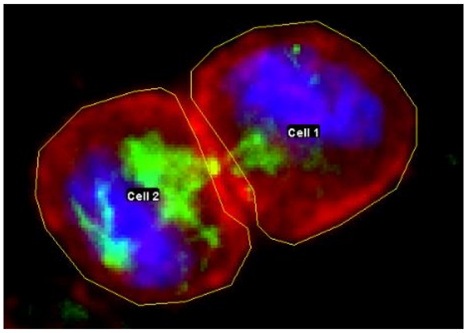


**Figure S7: Identification of mitotic cells and determination of regions of interest (ROI).**

Mitotic cells were selected based on nuclear (DAPI, blue staining) and β-tubulin (green staining) structures and imaged from late anaphase to telophase when daughter cells could be distinguished. The cell with higher intensities of CD8 (red staining) was defined as P1. The ROI was drawn around each cell excluding the contact area between the two cells. The ACD ratio (P1-P2)/(P1+P2) was calculated. A cell division was considered asymmetric when the CD8 intensity was 50% higher in one daughter cell compared to the other, leading to the threshold of 0.2.
